# IP3R2 mediated inter-organelle Ca^2+^ signaling orchestrates melanophagy

**DOI:** 10.1101/2025.03.27.645854

**Authors:** Suman Saurav, Rajender K Motiani

## Abstract

Organelle dynamics and crosstalk play a critical role in cellular functions thereby regulating physiological processes and pathological conditions. A variety of cellular processes are outcome of a balance between organelle biogenesis and degradation. Pigmentation is one such homeostatic state that is a result of melanosome biogenesis and melanosome degradation. Although melanosome biogenesis is partially understood, the melanosome degradation i.e. melanophagy remains largely unappreciated. Here, we reveal that Inositol 1,4,5-trisphosphate receptor 2 (IP_3_R2) is a negative regulator of melanophagy. In this study, we developed two *de novo* ratio metric imaging probes to study melanophagy in live-cells. Using these probes, biochemical assays, ultrastructural studies, confocal microscopy, molecular analyses and calcium imaging; we demonstrate that IP_3_R2, but not IP_3_R1 or IP_3_R3, keeps melanophagy in check. *In vivo* studies in zebrafish model system further substantiate IP_3_R2’s functional relevance in pigmentation. Mechanistically, IP_3_R2 silencing decreases mitochondrial Ca^2+^ uptake, augments ADP/ATP ratio and thereby activates melanophagy. Simultaneously, IP_3_R2 knockdown increases ER-lysosome proximity, enhances lysosomal Ca^2+^ levels and decreases lysosomal pH. This in turn activates lysosomal TRPML1 channel and stimulates nuclear translocation of TFEB transcription factor, which facilitates transcription of key autophagy and two known melanophagy drivers. Taken together, we uncover that IP_3_R2-mediated Ca^2+^ signaling across organelles is a critical determinant of melanophagy and thereby skin pigmentation. Hence, this signaling cascade offers potential therapeutic prospects for the management of pigmentary disorders and skin malignancies.

**Highlights:**

➢ IP_3_R2, but not IP_3_R1 and IP_3_R3, is a critical positive regulator of melanogenesis *in vitro* and *in vivo*.
➢ Generation and validation of two *de novo* ratio-metric live cells imaging probes reveal crucial role of IP_3_R2 in melanophagy.
➢ IP_3_R2 knockdown decreases mitochondrial Ca^2+^ uptake, augments ADP/ATP ratio and thereby activates melanophagy via AMPK-ULK1 pathway.
➢ IP_3_R2 silencing enhances ER-lysosomal proximity, elevates lysosomal Ca^2+^ levels and reduces lysosomal pH.
➢ IP_3_R2 knockdown stimulates lysosomal TRPML1 channel activity thereby facilitating nuclear translocation of TFEB transcription factor.
➢ TFEB transcriptionally upregulates genes involved in the melanophagy process, leading to enhanced degradation of melanosomes and decreased pigmentation.

## Introduction

Human skin pigmentation serves as a defense mechanism against harmful ultraviolet (UV) rays. Dysregulated pigmentation can lead to skin cancers and results in pigmentary disorders (Ahuja *et al*, 2025). The damaging effects of ultraviolet radiation are shielded by a natural photo protective pigment i.e. melanin. It is produced through the process of melanogenesis within pigment-producing cells called melanocytes (Natarajan *et al*, 2014a). Melanogenesis occurs in the specialized lysosome related organelle i.e. melanosomes (Hida *et al*, 2020; Ho & Ganesan, 2011). Skin pigmentation is an outcome of homeostatic balance between melanosome biogenesis, melanogenesis within melanosomes and melanosome degradation i.e. melanophagy. Although molecular mechanisms driving melanosome biogenesis are somewhat understood, the signaling cascades that drive melanophagy remain largely unappreciated.

Calcium (Ca^2+^) signaling is emerging as a critical regulator of pigmentation (Sharma *et al*, 2023; Ahuja *et al*, 2025; Bellono & Oancea, 2014). Ca^2+^ within melanocytes contributes to their function thereby regulating dendricity and pigmentation (Toyoda *et al*, 1999). Ca^2+^ helps in defining skin color by regulating tyrosinase activity, an enzyme critical for melanogenesis (BUFFEY *et al*, 1993). Further, UV rays activate G protein-coupled receptor (GPCR) in melanocytes and initiate Phospholipase C (PLC) signaling cascade. This in turn activates transient receptor potential ankyrin subtype 1 (TRPA1) and transient receptor potential channel vanilloid subtype 1 (TRPV1) channels leading to Ca^2+^ influx and induction of melanin synthesis (Jia *et al*, 2021; Bellono *et al*, 2013). Further, the UV induced Ca^2+^ influx in melanocytes stimulates melanosome transfer to keratinocytes which in turn provides photoprotection (Hu *et al*, 2017; Singh *et al*, 2017). While role of extracellular Ca^2+^ influx in regulating skin pigmentation is being appreciated, the role of organelle Ca^2+^ signaling is poorly understood.

We and others have recently demonstrated a crucial role of endoplasmic reticulum Ca^2+^ signaling in regulating skin pigmentation and controlling melanoma tumor progression (Cai *et al*, 2024; Tanwar *et al*, 2022b; Motiani *et al*, 2018). Further, mitochondrial matrix Ca^2+^ uptake via Mitochondrial Ca^2+^ Uniporter (MCU) complex drives skin pigmentation (Tanwar *et al*, 2024) by regulating melanosome biogenesis. On the other hand, outer mitochondrial membrane Ca^2+^ uptake channel, VDAC negatively regulates pigmentation by controlling transcription of genes involved in melanogenesis (Wang *et al*, 2022a). Interestingly, mitochondria tethers with melanosomes and that in turn regulates melanogenesis (Daniele *et al*, 2014; Tanwar *et al*, 2022a). Therefore, emerging literature suggest that organelle Ca^2+^ signaling can contribute to pigmentation. However, functional significance of Ca^2+^-driven inter-organelle crosstalk in regulating pigmentation is still unappreciated and that in melanophagy is completely unknown.

Here, we reveal that IP_3_R2 regulates Ca^2+^-driven ER-Mitochondria and ER-lysosomal crosstalk that in turn drives melanophagy to moderate pigmentation. Analysis of two independent microarray datasets showed that IP_3_R2 expression is directly associated with pigmentation levels. Targeted siRNA screening for IP_3_R isoforms identified IP_3_R2 as a negative regulator of pigmentation. Further, our overexpression and rescue experiments with wild-type and pore-dead IP_3_R2 mutant demonstrated that IP_3_R2 mediated ER Ca^2+^ release is essential for modulating pigmentation. Importantly, our *in vivo* experiments in zebrafish model recapitulated the *in vitro* observations at the organism level. Our robust mechanistic studies show that IP_3_R2 silencing enhances ER and lysosome proximity. This in turn augments lysosomal calcium levels and decreases lysosomal pH. It subsequently activates lysosomal TRPML1 channel and stimulates nuclear translocation of TFEB transcription factor. TFEB drives transcription of key autophagy and two known melanophagy drivers and thereby stimulates melanophagy. In summary, we have identified a novel signaling module that regulates melanophagy process and thereby modulate pigmentation. Since pigmentation protects from harmful UV radiations, this signaling cascade may offer potential therapeutic targets to manage pigmentary disorders and skin cancers.

## Results

### IP_3_R2, but not IP_3_R1 and IP_3_R3, positively regulates pigmentation

We have recently reported that physiological melanogenic stimuli α-melanocyte stimulating hormone (αMSH) generates IP_3_ thereby inducing release of Ca^2+^ from the IP_3_ receptors (IP_3_Rs) localized on the endoplasmic reticulum (Motiani *et al*, 2018; Tanwar *et al*, 2024). However, the functional relevance of IP_3_Rs in the pigmentation remains completely unknown. We had earlier performed two independent microarrays to identify novel regulators of pigmentation (Natarajan *et al*, 2014b; Motiani *et al*, 2018). The first microarray was performed on B16 mouse melanoma cells while they autonomusly acquired pigmentation in 6-7 days upon low density (LD) cultutring (**Fig 1A**). We examined levels of IP_3_Rs in the microarrays and found that mRNA expression of IP_3_Rs is augmented with the increase in pigmentation. This observation was further corroborated through quantitative real-time PCR (qRT-PCR) analysis, which demonstrated that the IP_3_R2 isoform exhibited a more substantial increase in expression than the other isoforms (**Fig 1B**). The second microarray was performed on primary human melanocytes, which were chemically stimulated either with tyrosine, a pigment inducing agent or phenylthiourea (PTU), a de-pigmentary agent (**Fig 1C**). In this microarray only the expression of IP_3_R2 was directly proportional to pigmentation levels. We corroborated this observation by performing qRT-PCRs and found that upon tyrosine mediated hyperpigmentation IP_3_R2 expression was enhanced (**Fig 1D**) while upon PTU driven hypopigmentation IP_3_R2 expression is decreased (**Fig 1E**).

**Figure 1:**
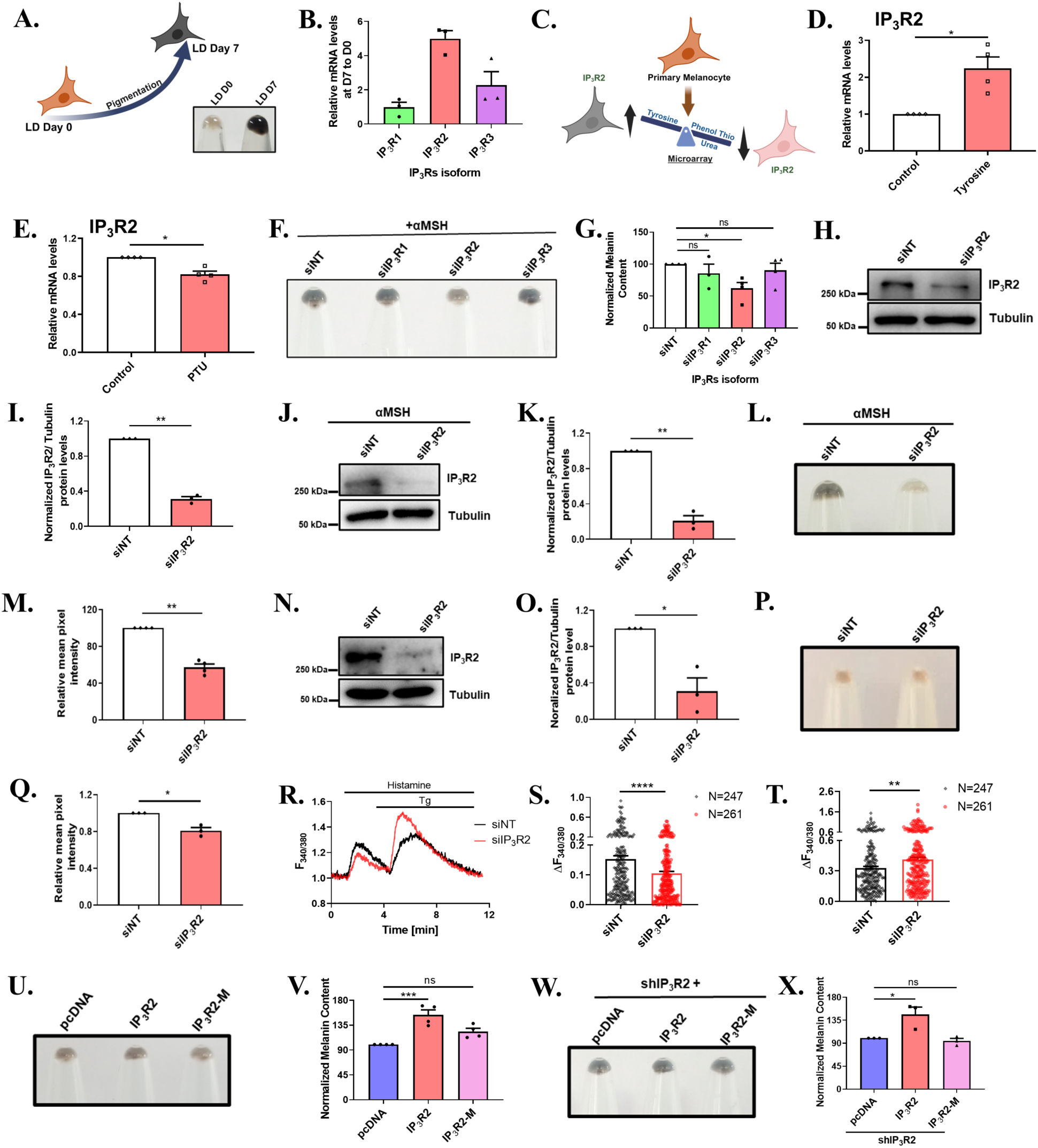
IP_3_R2 is a positive regulator of pigmentation. (A) Pictorial representation and cell pellet image demonstrating increased pigmentation of B16 cells over seven days, from day0 to day7. (B) qRT PCR analysis showing quantification of IP_3_R1, IP_3_R2, and IP_3_R3 mRNA in B16 cells (N=3). (C) Microarray analysis of IP_3_R2 in primary human melanocyte showing pictorial representation of microarray stimulated with L-tyrosine and Phenylthiourea. (D) qRT PCR analysis showing increase in IP_3_R2 mRNA expression after 72hrs of treatment with 1mM L-Tyrosine (N=4). (E) qRT PCR analysis showing decrease in IP_3_R2 mRNA expression after 72hrs of treatment with 200µM Phenylthiourea (N=4). (F) Representative image of pellet pictures demonstrates B16 cells of LD day6, transfected with either non-targeted siRNA or siRNA specifically targeting IP_3_ receptor type1, type2, and type3 on LD day3. (G) Bar graph showing melanin content estimation of B16 cells harvested 72hrs post siRNA transfection at LD day6 (N=4). (H) Representative image of immunoblot showing expression of IP_3_R2 in B16 cells after siRNA silencing of IP_3_R2 in LD model system. (I) Bar graph showing the densitometry of IP_3_R2 band normalized to β-Tubulin (N=3). (J) Representative image of immunoblot showing the expression of IP_3_R2 in B16 cells after 72hrs of siRNA silencing of IP_3_R2, with a 48hrs exposure to 1µM of αMSH. (K) Bar graph showing the densitometry of the IP_3_R2 band normalized to β-Tubulin (N=3). (L) Representative image of pellet pictures shows B16 cells 72hrs after transfection with either non-targeted siRNA or siRNA specifically targeting IP_3_R2, along with a 48hrs exposure to 1µM αMSH. (M) Bar graph showing mean pixel intensity of B16 cells harvested 72hrs post siRNA transfection in addition to 48hrs exposure to 1µM αMSH (N=4). (N) Representative image of immunoblot showing the expression of IP_3_R2 in lightly pigmented primary human melanocytes after 72hrs of IP_3_R2 silencing. (O) Bar graph showing the densitometry of the IP_3_R2 band normalized to β-Tubulin (N=3). (P) Representative image of pellet picture showing primary human melanocytes 72hrs post transfection with either non-targeted siRNA or siRNA specifically targeting IP_3_R2. (Q) Bar graph showing mean pixel intensity of lightly pigmented primary human melanocytes harvested 72hrs post-siRNA transfection (N=3). (R) Representative traces of Fura2AM based Ca^2+^ imaging in B16 cells stimulated with the Histamine followed by Tg after 72hrs of siRNA transfection. (S) Bar graph showing the quantification of Ca^2+^ imaging traces stimulated with 50µM histamine, where ‘N’ denotes the total number of cells imaged. (T) Bar graph showing the quantification of Ca^2+^ imaging traces stimulated with 2µM Tg, where ‘N’ denotes the total number of cells imaged. (U) Representative image of pellet picture showing B16 cells transfected with IP_3_R2 and IP_3_R2-M in the LD model system. (V) Bar graph showing the melanin content in B16 cells transfected with IP_3_R2 and IP_3_R2-M (N=4). (W) Representative image of pellet picture showing B16 cells overexpressed with IP_3_R2 and IP_3_R2-M in stable IP_3_R2 knockdown background. (X) Bar graph showing the melanin content in B16 cells overexpressed with IP_3_R2 and IP_3_R2-M in stable IP_3_R2 knockdown background (N=3). Data presented are mean ± SEM. For statistical analysis, Dunnett’s multiple comparisons test was performed for panels G, V, X, while one sample t-test was performed for panels D, E, I, K, M, O, Q and an Unpaired t-test was performed for panels S, T using GraphPad Prism software. Here, ‘ns’ means non-significant; * p <0.05, ** p < 0.01 and *** p < 0.001.

Collectively, the microarrays and qRT-PCR validation show that the levels of IP_3_Rs, in particular IP_3_R2, are positively associated with pigmentation levels in two independent cellular models. To investigate the role of the IP_3_R isoforms in melanogenesis, we carried out a targeted small interfering RNA (siRNA) screen against the three IP_3_R isoforms in B16 cells. We first validated the efficacy of siRNAs by performing qRT-PCRs. This analysis revealed a drastic reduction in the mRNA levels of the targeted IP_3_R isoform, while the expression of other isoforms remained unchanged or showed a minor change (**Supplementary Fig 1A-C**). Next, we examined the effect of IP_3_R isoforms silencing on pigmentation in B16 LD pigmentation model. We observed a phenotypic decrease in the pigmentation upon IP_3_R2 silencing while knockdown of IP_3_R1 and IP_3_R3 showed no phenotypic changes as compared to control siNon-targeting (siNT) condition (**Fig 1F**). We further validated these phenotypic observations by performing quantitative melanin content assays (**Fig 1G**).

Since we observed changes in pigmentation upon only IP_3_R2 silencing, we directed our efforts in precisely delineating the role of IP_3_R2 in pigmentation. First of all, we confirmed IP_3_R2 silencing at protein-level with both siRNA mediated transient and shRNA driven stable knockdown of IP_3_R2 in B16 cells. We observed a significant reduction in IP_3_R2 expression in both these conditions (**Fig 1H-I and Supplementary Fig 1D-E**). We subsequently analyzed phenotypic and quantitative change in pigmentation upon stable IP_3_R2 silencing. As expected, we found a significant decrease in pigmentation (**Supplementary Fig 1F-G**). Next, we examined role of IP_3_R2 in αMSH stimulated physiological pigmentation. We observed a robust decrease in αMSH induced pigmentation upon both transient (**Fig 1J-M**) and stable silencing of IP_3_R2 in B16 cells (**Supplementary Fig 1H-K**). Finally, we validated critical role of IP_3_R2 in human skin pigmentation by performing experiments in primary human melanocytes. IP_3_R2 silencing with a human IP_3_R2 siRNA led to a significant decrease in both IP_3_R2 protein expression and the pigmentation phenotype as compared to the control non-targeting siRNA (**Fig 1N-Q**).

Since IP_3_R2 is a key ER Ca^2+^ release channel, we next examined the effect of IP_3_R2 knockdown on histamine (inducer of Ca^2+^ release via IP_3_Rs) stimulated ER Ca^2+^ release and Thapsigargin (Tg) mediated bulk ER Ca^2+^ mobilization. Our Fura2AM based live-cell Ca^2+^ imaging experiments show that upon stimulation with histamine the IP_3_R2 knockdown condition exhibited reduced Ca^2+^ release as compared to control siRNA (**Fig 1R-S**). Consequently, the Tg mediated bulk ER Ca^2+^ mobilization was higher in siIP_3_R2 condition in comparison to control siRNA (**Fig 1R, 1T**). This indicates that IP_3_R2 plays a functional role in mediating ER Ca^2+^ release in melanocytes. To investigate the significance of IP_3_R2 mediated ER Ca^2+^ release in regulating pigmentation we utilized a non-conductive pore mutant of IP_3_R2 (IP_3_R2-M), which cannot facilitate Ca^2+^ release and a wild-type functional IP_3_R2 construct (Bartok *et al*, 2019). Using quantitative qRT-PCR, we evaluated the expression levels of IP_3_R2 and IP_3_R2-M. Our analysis revealed a significant and specific up regulation of the IP_3_R2 isoform with wild type IP_3_R2 (**Supplementary Fig 1L**) and IP_3_R2-M construct (**Supplementary Fig 1M**). Next, we examined the effect of functional IP_3_R2 and IP_3_R2-M overexpression on LD pigmentation. We observed that overexpression of the functional IP_3_R2 isoform resulted in increased pigmentation, while the overexpression of IP_3_R2-M did not enhanced phenotypic pigmentation (**Fig 1U**). We further quantitated the differences via melanin content assays, which corroborated the qualitative images (**Fig 1V**). We further validated these observations by rescuing the functional IP3R2 and IP3R2-M expression in the stable IP_3_R2 knockdown background. Both the phenotypic and quantitative analysis of pigmentation clearly demonstrated that the overexpression with the functional IP_3_R2 isoform recovered the pigmentation phenotype (**Fig 1W-X**). However, the expression of the IP_3_R2-M failed to enhance the pigmentation phenotype (**Fig 1W-X**). This data highlights that the Ca^2+^ release via IP_3_R2 is required for driving melanin production. Taken together, our data from four independent *in vitro* model systems (LD pigmentation in B16 cells, αMSH-stimulated physiological melanogenesis, pigmentation in primary human melanocytes and overexpression/rescue experiments) elegantly demonstrate that IP_3_R2 is a positive regulator of pigmentation.

### IP_3_R2 regulates pigmentation *in vivo*

To examine the role of IP_3_R2 in pigmentation *in vivo*, we performed IP_3_R2 loss of function and gain of function studies in zebrafish model system. Zebrafish is a well-established model organism for pigmentation studies (Motiani *et al*, 2018; Tanwar *et al*, 2024; Qu *et al*, 2023). We carried out IP_3_R2 loss of function studies by performing IP_3_R2 knockdown by injecting morpholinos targeting IP_3_R2 at single cell stage zebrafish embryos. The qRT-PCR analysis demonstrated a significant decrease in IP_3_R2 expression upon IP_3_R2 morpholino injections compared to the scrambled morpholino (**Fig 2A**). Microscopic examination revealed that IP_3_R2 silencing results in reduction of pigmentation in zebrafish embryos compared to the control group at 48 hours post-fertilization (**Fig 2B**). We further validated the phenotypic observation by performing melanin content assays and observed around 35% reduction in pigmentation upon IP_3_R2 knockdown (**Fig 2C**).

**Figure 2:**
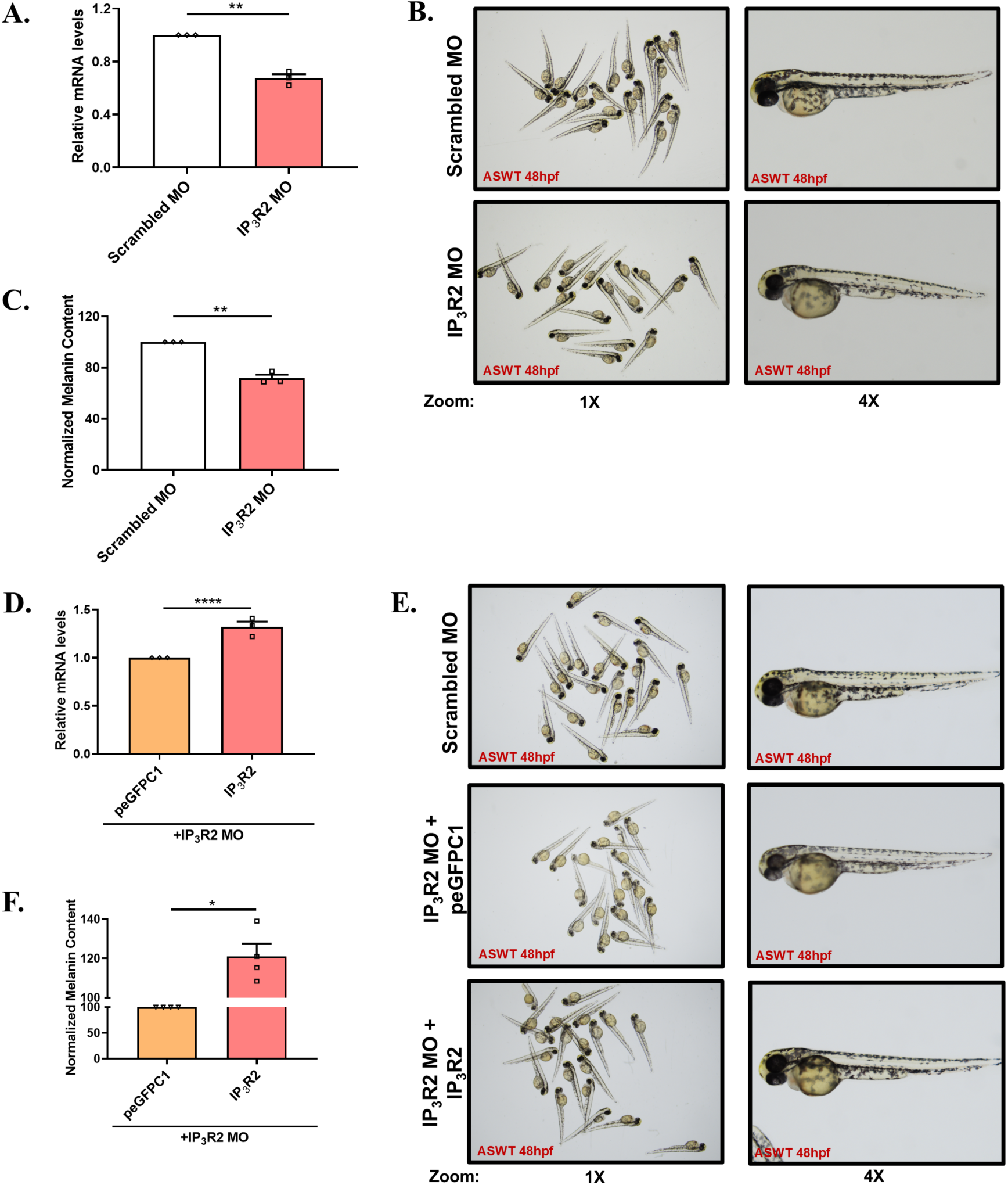
IP_3_R2 drives pigmentation *in vivo*. (A) qRT PCR analysis showing decrease in IP_3_R2 mRNA expression after 48hrs post fertilization (hpf) on zebrafish larvae injected with 400µM IP_3_R2 morpholino (N=3). (B) Representative images of Assam wild type (ASWT) zebrafish after 48hpf injected with 400µM IP_3_R2 morpholino. (C) Bar graph showing melanin content estimation of zebrafish after 48hpf injected with 400µM IP_3_R2 morpholino (N=3 with 20 embryos in each “N”). (D) qRT PCR analysis showing increase in IP_3_R2 mRNA expression after 48hpf on zebrafish larvae injected with 400µM IP_3_R2 morpholino and rescued with 120ng of IP_3_R2 mRNA (N=4). (E) Representative images of ASWT zebrafish embryos after 48hpf injected with 400µM IP_3_R2 morpholino and rescued with 120ng of IP_3_R2 mRNA. (F) Bar graph showing melanin content estimation of zebrafish embryos after 48hpf injected with 400µM IP_3_R2 morpholino and rescued with 120ng of IP_3_R2 mRNA (N=3 with 20 embryos in each “N”). Data presented are mean ± SEM. For statistical analysis, one sample t-test was performed for panels A, C, D, F using GraphPad Prism software. Here, * p <0.05, ** p < 0.01 and **** p < 0.0001.

We next conducted rescue experiments to corroborate the effects observed upon IP3R2 silencing. We co-injected either IP_3_R2 or control (GFP) mRNA into zebrafish embryos along with IP_3_R2 morpholinos. qRT-PCR demonstrated that the embryos receiving IP_3_R2 mRNA exhibited significantly higher IP_3_R2 expression levels in comparison to the control group (**Fig 2D**). Importantly, the embryos rescued with IP_3_R2 mRNA displayed an increase in pigmentation phenotype, which we further substantiated via melanin content assay (**Fig 2E-F**). This data clearly demonstrates that IP_3_R2 regulates pigmentation *in vivo*. Collectively, our comprehensive *in vitro* and *in vivo* studies establish IP_3_R2 as a novel positive regulator of pigmentation.

### IP_3_R2 knockdown enhances stability of melanogenic proteins

To further substantiate the phenotypic observations, we analyzed the mRNA expression of critical melanogenic genes following IP_3_R2 knockdown. We first confirmed IP_3_R2 silencing temporally by performing qRT-PCR (**Fig 3A**). Subsequently, we assessed the mRNA levels of the melanosomal structural protein (Pre-melanosome Protein 17, i.e., PMEL17 or GP100) and the key melanogenic enzymes (Tyrosinase and DCT). Interestingly, we did not observe significant changes in the mRNA expression of these melanogenic genes (**Fig 3B-D**). We next performed western blotting to investigate the impact of IP_3_R2 knockdown on the protein expression of these melanogenesis regulators. Interestingly, despite the decrease in pigmentation upon IP_3_R2 silencing, we observed a significant increase in the melanogenic proteins (**Fig 3E-J**). Collectively, this suggests that the decrease in pigmentation observed upon IP_3_R2 silencing is not due to transcriptional regulation of melanogenic genes but may involve post-translational mechanisms or alterations in protein stability.

**Figure 3:**
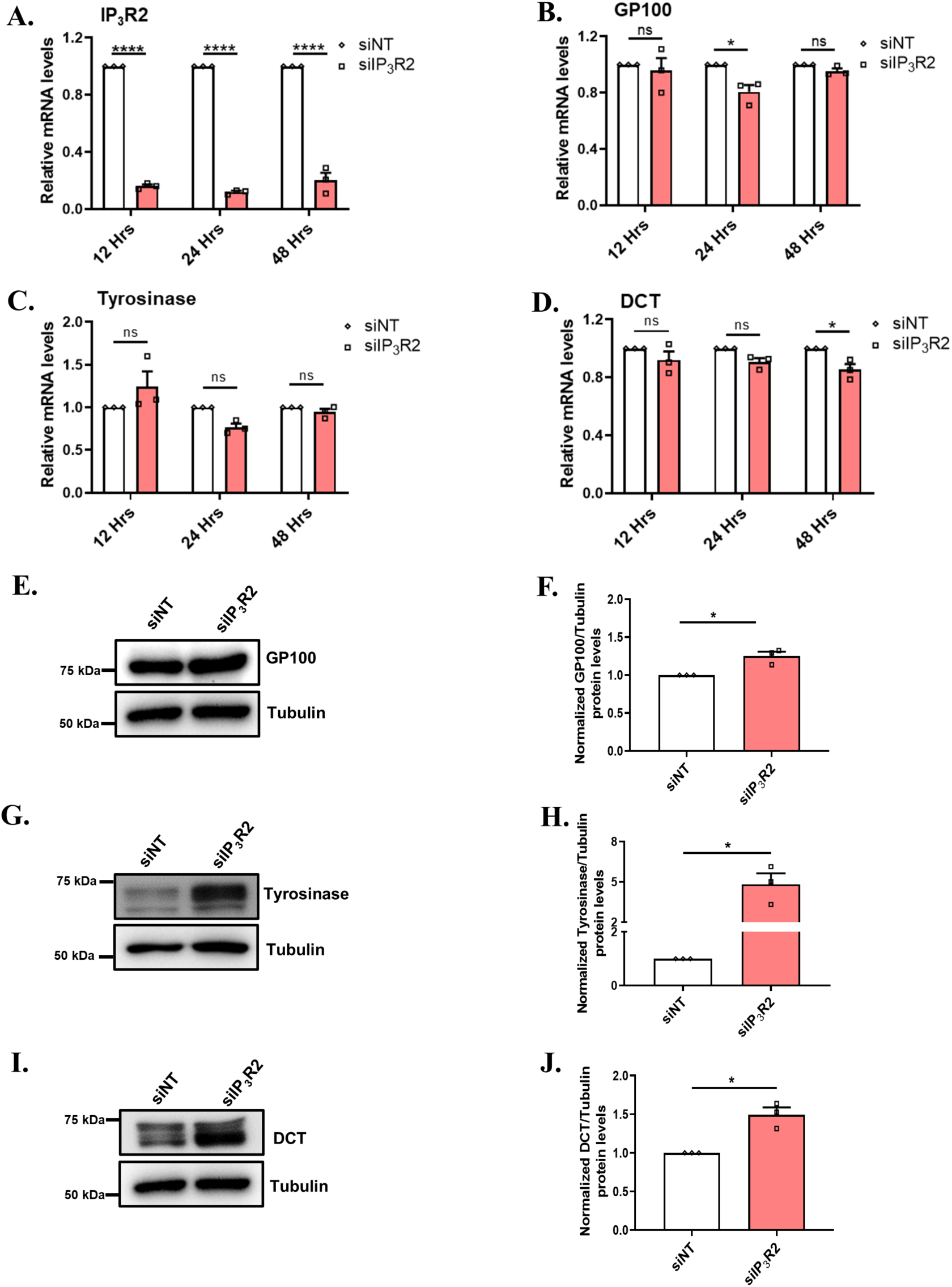
IP_3_R2 knockdown increases the stability of melanogenic proteins. (A) qRT PCR analysis showing expression of IP_3_R2 in B16 cells incubated with 1µM αMSH for 12, 24, and 48hrs following 24hrs of transfection with siNT or siIP_3_R2 (N=3). (B) qRT PCR analysis showing expression of GP100 in B16 cells incubated with 1µM αMSH for 12, 24, and 48hrs following 24hrs of transfection with siNT or siIP_3_R2 (N=3). (C) qRT PCR analysis showing expression of Tyrosinase in B16 cells incubated with 1µM αMSH for 12, 24, and 48hrs following 24hrs of transfection with siNT or siIP_3_R2 (N=3). (D) qRT PCR analysis showing expression of DCT in B16 cells incubated with 1µM αMSH for 12, 24, and 48hrs following 24hrs of transfection with siNT or siIP_3_R2 (N=3). (E) Representative image of immunoblot showing expression of GP100 in B16 cells after siRNA silencing of IP_3_R2 in LD model system. (F) Bar graph showing the densitometry of GP100 band normalized to β-Tubulin (N=3). (G) Representative image of immunoblot showing expression of Tyrosinase in B16 cells after siRNA silencing of IP_3_R2 in LD model system. (H) Bar graph showing the densitometry of Tyrosinase band normalized to β-Tubulin (N=3). (I) Representative image of immunoblot showing expression of DCT in B16 cells after siRNA silencing of IP_3_R2 in LD model system. (J) Bar graph showing the densitometry of DCT band normalized to β-Tubulin (N=3). Data presented are mean ± SEM. For statistical analysis, Sidak’s multiple comparisons test was performed for panels A, B, C, D, and one sample t-test was performed for panels F, H, J using GraphPad Prism software. Here, ‘ns’ means non-significant; * p <0.05, and **** p < 0.0001.

### IP_3_R2 silencing enhances melanogenic proteins’ half-life and induces autophagy flux

We next directed our efforts to determine the underlying mechanism through which IP_3_R2 downregulation increases melanogenic proteins expression. We investigated protein stability of the key melanogenic regulators in presence of protein synthesis inhibitor cycloheximide. Our data show that IP_3_R2 levels are decreased in IP_3_R2 silenced condition (**Fig 4A-B**), while the melanogenic proteins DCT, Tyrosinase and GP100, exhibited higher stability in the IP_3_R2 knockdown condition (**Fig4A and 4C-E**). Earlier studies have demonstrated that impairment of protein degradation stimulates autophagy as a cytoprotective mechanism (Williams *et al*, 2013; Li *et al*, 2019; Su & Wang, 2020; Pan *et al*, 2020). Therefore, we studied autophagy induction upon IP_3_R2 knockdown. We observed that IP_3_R2 silencing leads to an increase in LC3II levels in both LD pigmentation (**Fig 4F-G**) and αMSH induced pigmentation (**Fig 4H-I**) models. These results suggest that IP_3_R2 knockdown increases melanogenic proteins stability, which in turn activates autophagy. To substantiate these observations, we utilized the pMRX-IP-GFP-LC3-RFP reporter system, a powerful experimental approach that enables the quantification of autophagic flux by calculating the ratio of GFP to RFP signals (Kaizuka *et al*, 2016). IP_3_R2 silencing significantly reduced GFP/RFP ratio of the GFP-LC3-RFP reporter system in comparison to control conditions thereby indicating enhanced autophagic flux upon IP_3_R2 knockdown (**Fig 4J-K**). We further corroborated this by assessing autophagic flux using bafilomycin, an inhibitor of autophagosome-lysosome fusion. In the presence of bafilomycin, the IP_3_R2 knockdown cells showed a significantly increased LC3-II protein levels suggesting higher accumulation of autophagosomes, as compared to bafilomycin-treated control cells (**Fig 4L-M**). Taken together, these biochemical and live cell microscopy data clearly demonstrate that IP_3_R2 silencing enhances autophagic flux in melanocytes.

**Figure 4:**
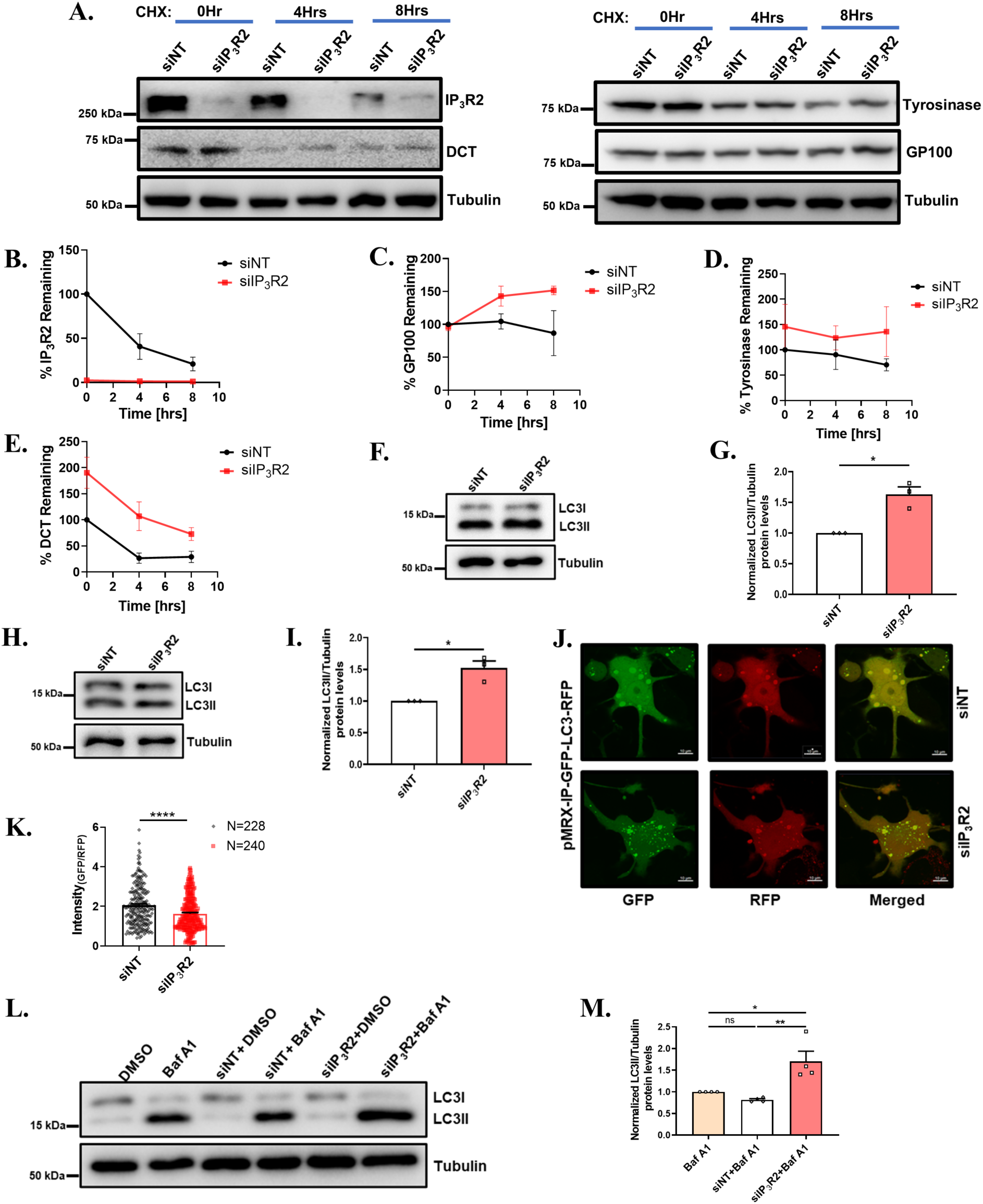
IP_3_R2 silencing stabilize melanogenic proteins and triggers autophagy. (A) Representative image of immunoblot showing expression of the IP_3_R2 and the melanogenic enzymes DCT, Tyrosinase, and DCT in B16 cells for different time interval of cycloheximide treatment after siRNA transfections. (B) The line graph shows the percentage of the proteins remaining relative to 0hr of cycloheximide treatment by densitometry of the IP_3_R2 band normalized to β-Tubulin. (C) The line graph shows the percentage of the proteins remaining relative to 0hr of cycloheximide treatment by densitometry of the GP100 band normalized to β-Tubulin. (D) The line graph shows the percentage of the proteins remaining relative to 0hr of cycloheximide treatment by densitometry of the Tyrosinase band normalized to β-Tubulin. (E) The line graph shows the percentage of the proteins remaining relative to 0hr of cycloheximide treatment by densitometry of the DCT band normalized to β-Tubulin. (F) Representative image of immunoblot showing expression of LC3II in B16 cells after siRNA silencing of IP_3_R2 in LD model system. (G) Bar graph showing the densitometry of the IP_3_R2 band normalized to β-Tubulin (N=3). (H) Representative image of immunoblot showing expression of LC3II in B16 cells after siRNA silencing of IP_3_R2. (I) Bar graph showing the densitometry of the IP_3_R2 band normalized to β-Tubulin (N=3). (J) Representative images of confocal imaging in B16 cells after transfected with siNT or siIP_3_R2 along with pMRX-IP-GFP-LC3-RFP probe, scale bar, 10µm. (K) Bar graph shows the quantification of the autolysosome intensity in the GFP:RFP ratio, where ‘N’ denotes the number of cells. (L) Representative image of immunoblot showing expression of LC3II in B16 cells after siRNA transfection including 6hrs of treatment with 100nM of the lysosomal inhibitor Bafilomycin A1. (M) Bar graph showing the densitometry of the LC3II band normalized to β-Tubulin (N=4). Data presented are mean ± SEM. For statistical analysis, one sample t-test was performed for panels G, I, an unpaired t-test was performed for panel K, and Dunnett’s multiple comparisons test was performed for panel M using GraphPad Prism software. Here, ‘ns’ means non-significant; * p <0.05, ** p < 0.01, and **** p < 0.0001.

### IP_3_R2 silencing induces melanosome degradation

We observed an impaired proteasomal degradation of critical melanogenic proteins localized on melanosomes in the IP_3_R2 knockdown condition. This may increase melanophagy, the selective autophagic degradation of melanosomes, as a cytoprotective mechanism to maintain protein homeostasis (Zhu *et al*, 2010; Xiong *et al*, 2023). Therefore, we developed two *de novo* ratiometric fluorescent probes to study melanophagy flux in live cells. In the first probe, we tagged Tyrosinase, a protein exclusively localized on melanosomes, with mCherry and eGFP (**Fig 5A**). The ratiometric, mCherry-Tyrosinase-eGFP probe, functions based on the differential stability of the green and red fluorescent proteins in low pH milieu. The acidic environment within the lysosome (pH <5.2) selectively quenches the fluorescent signal of eGFP and has minimal impact on the mCherry signal (Park *et al*, 2020). We validated the functionality of mCherry-Tyrosinase-eGFP probe by treating B16 cells expressing this probe with rapamycin, a known autophagy inducer, which resulted in a significant decrease in the GFP signal and a lower GFP/RFP ratio than the control condition (**Supplementary Fig 2A**). Conversely, treatment with bafilomycin A1, an inhibitor of autophagosome-lysosome fusion, led to an increase in the GFP signal and a higher GFP/RFP ratio relative to the control (**Supplementary Fig 2B**). These assays exhibit that mCherry-Tyrosinase-eGFP probe can effectively measure the quantitative changes in autophagic flux. We next used this probe to study melanophagy flux upon IP_3_R2 silencing, we observed a significant decrease GFP signal and a lower GFP/RFP ratio in the IP_3_R2 knockdown condition compared to the control non-targeting siRNA condition (**Fig 5B-C**), indicating enhanced melanophagy upon IP_3_R2 silencing.

**Figure 5:**
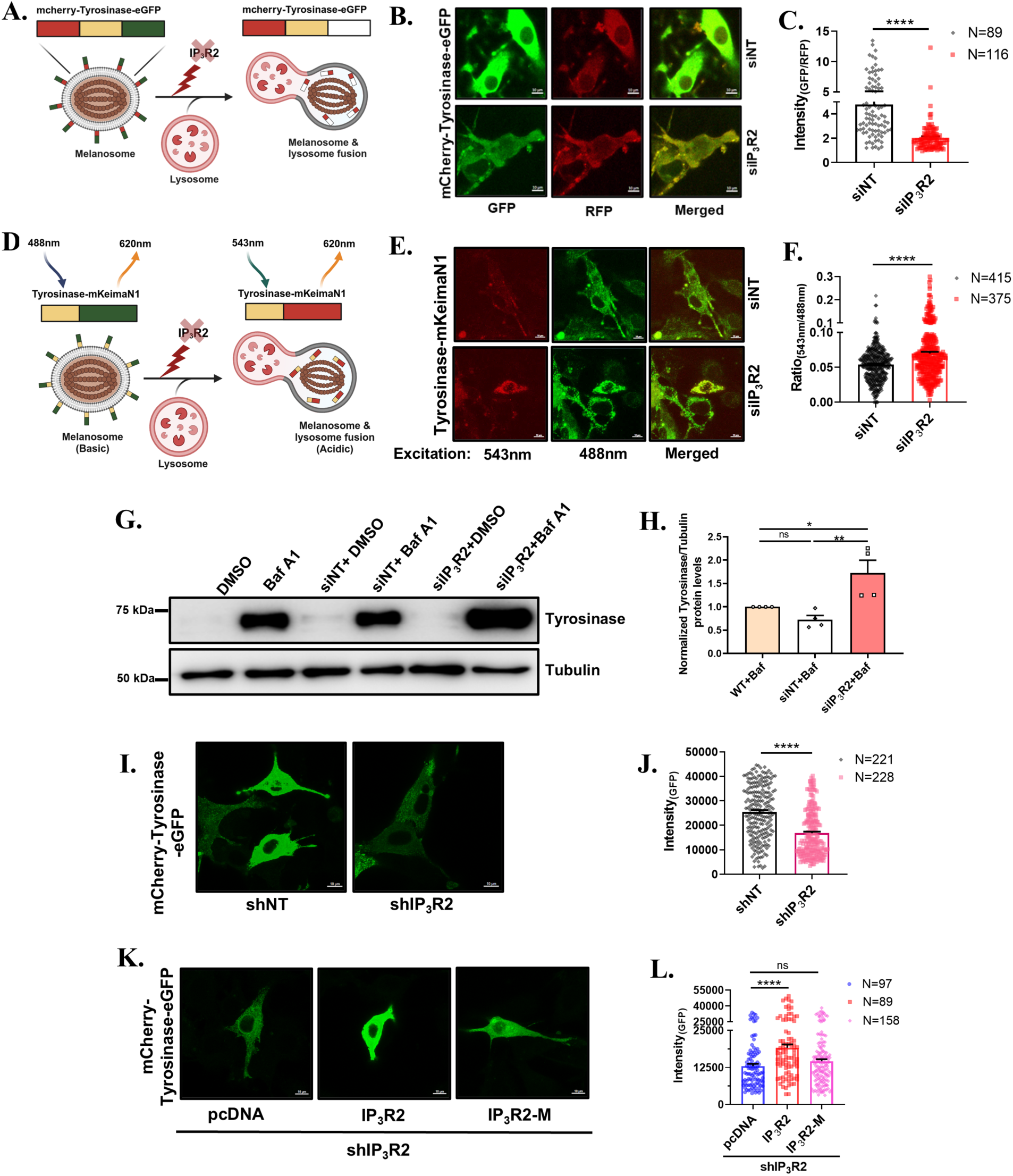
IP_3_R2 silencing enhances melanophagy. (A) A schematic illustration demonstrating the melanophagy flux upon IP_3_R2 silencing by utilizing a mCherry-Tyrosinase-EGFP construct. (B) Representative images of confocal imaging in B16 cells after transfected with siNT or siIP_3_R2 along with mCherry-Tyrosinase-EGFP probe, scale bar, 10µm. (C) Bar graph shows the quantification of the autolysosome intensity in the GFP:RFP ratio, where ‘N’ denotes the number of cells. (D) A schematic illustration demonstrating the melanophagy flux upon IP_3_R2 silencing by utilizing Tyrosinase-mKeimaN1 construct. (E) Representative images of confocal imaging in B16 cells after transfected with siNT or siIP_3_R2 along with Tyrosinase-mKeimaN1 probe, scale bar, 10µm. (F) Bar graph shows the quantification of the autolysosome intensity in the 543nm:488nm ratio, which is alternatively excited at 543nm and 488nm, where ‘N’ denotes the number of cells. (G) Representative image of immunoblot showing expression of Tyrosinase in B16 cells after siRNA transfection including 6hrs of treatment with 100nM of the lysosomal inhibitor Bafilomycin A1. (H) Bar graph showing the densitometry of the Tyrosinase band normalized to β-Tubulin (N=4). (I) Representative images of confocal imaging in B16 cells having stable IP_3_R2 knockdown transfected with mCherry-Tyrosinase-EGFP construct, scale bar, 10µm. (J) Bar graph showing the quantification of the GFP intensity (mCherry inherently present in the shRNA), where ‘N’ denotes the number of cells imaged. (K) Representative images of confocal imaging in B16 cells having stable IP_3_R2 knockdown demonstrating rescue with IP_3_R2 and IP_3_R2-M, scale bar, 10µm. (L) Bar graph showing the quantification of the GFP intensity (mCherry inherently present in the shRNA), where ‘N’ denotes the number of cells imaged. Data presented are mean ± SEM. For statistical analysis, an unpaired t-test was performed for panels C, F, and J Dunnett’s multiple comparisons test was performed for panel H and L using GraphPad Prism software. Here, ‘ns’ means non-significant; * p <0.05, ** p < 0.01, and **** p < 0.0001.

To further substantiate this observation, we generated an additional fluorescent ratiometric probe using mKeima. mKeima is a pH-sensitive fluorescent protein derived from coral with emission spectra peaking at 620nm and a biphasic excitation spectra peaking at 543nm (Red) and 488nm (Green) in acidic and neutral environments respectively (Katayama *et al*, 2011; Yoshii & Mizushima, 2017). We developed a ratiometric fluorescent probe by inserting the tyrosinase protein upstream of the mKeimaN1. This fluorescent reporter system enables the monitoring of melanosomal degradation within the acidic lysosomal environment. The signal of the tyrosinase-mKeimaN1 fusion protein increases at 543nm excitation in response to the low pH conditions resulting in an increased 543nm/488nm signal ratio (**Fig 5D**). The functionality of the tyrosinase-mKeimaN1 probe was assessed by treating tyrosinase-mKeimaN1 transfected B16 cells with rapamycin and bafilomycin A1. The rapamycin treatment increased red (543nm excitation) fluorescence signal resulting in a higher 543/488nm signal ratio while bafilomycin A1 treatment resulted in a diminished red (543nm excitation) fluorescence signal and a lower 543/488nm signal ratio (**Supplementary Fig 2C-D**). This data elegantly demonstrates that the ratiometric tyrosinase-mKeimaN1 probe can effectively monitor changes in melanophagy. Utilizing the tyrosinase-mKeimaN1 probe, we observed a significant increase in the red (543nm excitation) fluorescence signal and a higher 543/488nm signal ratio in IP_3_R2 knockdown condition in comparison to the control siRNA group (**Fig 5E-F**).

To further corroborate these microscopic observations, we performed biochemical assays to study melanophagy flux upon IP_3_R2 silencing. We employed bafilomycin, an inhibitor of autophagosome-lysosome fusion, to study the accumulation of melanosomal proteins upon IP_3_R2 knockdown. In the presence of bafilomycin, IP_3_R2 silenced cells exhibited a more pronounced accumulation of melanosomes, as indicated by elevated tyrosinase levels as compared to bafilomycin-treated control non-targeting siRNA (siNT) condition (**Fig 5G-H**) highlighting that IP_3_R2 knockdown increases melanophagy flux. We next performed ultrastructural studies to further substantiate the increase in melanophagy upon IP_3_R2 silencing. Our Transmission Electron Microscopy (TEM) data demonstrates a significant decrease in the melanosome numbers in IP_3_R2 knockdown cells in comparison to the control group **(Supplementary Fig 3A-B)**.

Finally, we validated IP_3_R2’s role in melanophagy by performing rescue experiments in cells with stable IP_3_R2 knockdown. We examined melanophagy using mCherry-Tyrosinase-eGFP probe in control cells and in the cells with stable IP_3_R2 knockdown. Since shRNA targeting IP_3_R2 was tagged with RFP, we focused on GFP fluorescence intensity as readout of decrease in melanophagy (**Fig 5I-J**). Further, we performed rescue experiments with either wild-type IP_3_R2 or IP_3_R2-M and corresponding empty vector control in stable IP_3_R2 knockdown cells (**Fig 5K-L**). Our data show that stable IP_3_R2 knockdown enhances melanophagy (**Fig 5I-J**) which could be rescued with wild-type IP_3_R2 but not IP_3_R2-M (**Fig 5K-L**). To corroborate these observations, we performed ultrastructural studies with the wild-type IP_3_R2 and IP_3_R2-M rescue experiments in stable IP_3_R2 knockdown condition. The TEM data shows a significant increase in melanosome number in cells rescued with IP_3_R2 while rescue with IP_3_R2 M shows similar melanosome number as compared to stable IP_3_R2 knockdown **(Supplementary Fig 3C-D)**. These findings demonstrate that the Ca^2+^ release function of IP_3_R2 is a critical determinant of melanophagy. Collectively, the confocal imaging with two ratiometric probes, biochemical assays, ultrastructural studies, and the rescue experiments reveal that IP_3_R2 negatively regulates melanophagy.

### IP_3_R2 knockdown impairs mitochondrial Ca^2+^ uptake and triggers melanophagy

To elucidate the underlying mechanism by which IP_3_R2 knockdown triggers melanophagy, we investigated the potential impact of IP_3_R2 silencing on mitochondrial Ca^2+^ dynamics. Mitochondrial Ca^2+^ uptake is a critical determinant of cellular activities (Pathak & Trebak, 2018; Tanwar *et al*, 2020; Mammucari *et al*, 2018). Decrease in IP_3_R expression and/or function diminishes the IP_3_R mediated Ca^2+^ signals thereby hampering mitochondrial function and reducing ATP generation (Fink *et al*, 2017). We measured mitochondrial Ca^2+^ uptake, using the genetically encoded fluorescent Ca^2+^ indicator probe Cepia2mt, in response to histamine (an IP_3_-generating stimuli) in control siNT and siIP_3_R2 transfected cells. We observed that IP_3_R2 knockdown resulted in a substantial decrease in the mitochondrial Ca^2+^ uptake (**Fig 6A-B)**. We substantiated these findings by utilizing a B16 cell line with stable IP_3_R2 knockdown and a corresponding control cell line. We assessed histamine induced mitochondrial Ca^2+^ uptake in these cells and validated that IP_3_R2 depletion leads to reduction in mitochondrial Ca^2+^ influx (**Fig 6C-D**). To further corroborate the role of IP_3_R2 function in regulating mitochondrial Ca^2+^ dynamics, we performed rescue experiments in the stable IP_3_R2 knockdown cells. Either the full-length IP_3_R2 and IP_3_R2-M, which cannot facilitate Ca^2+^ release, were ectopically expressed in the stable IP_3_R2 knockdown cells. Our results show that overexpression of the wild-type IP_3_R2 restored the mitochondrial Ca^2+^ uptake while the IP_3_R2-M failed to rescue the mitochondrial Ca^2+^ uptake in stable IP_3_R2 knockdown cells (**Fig 6E-F**). These findings suggest that the Ca^2+^ release activity of IP_3_R2 plays a critical role in facilitating mitochondrial Ca^2+^ uptake.

**Figure 6:**
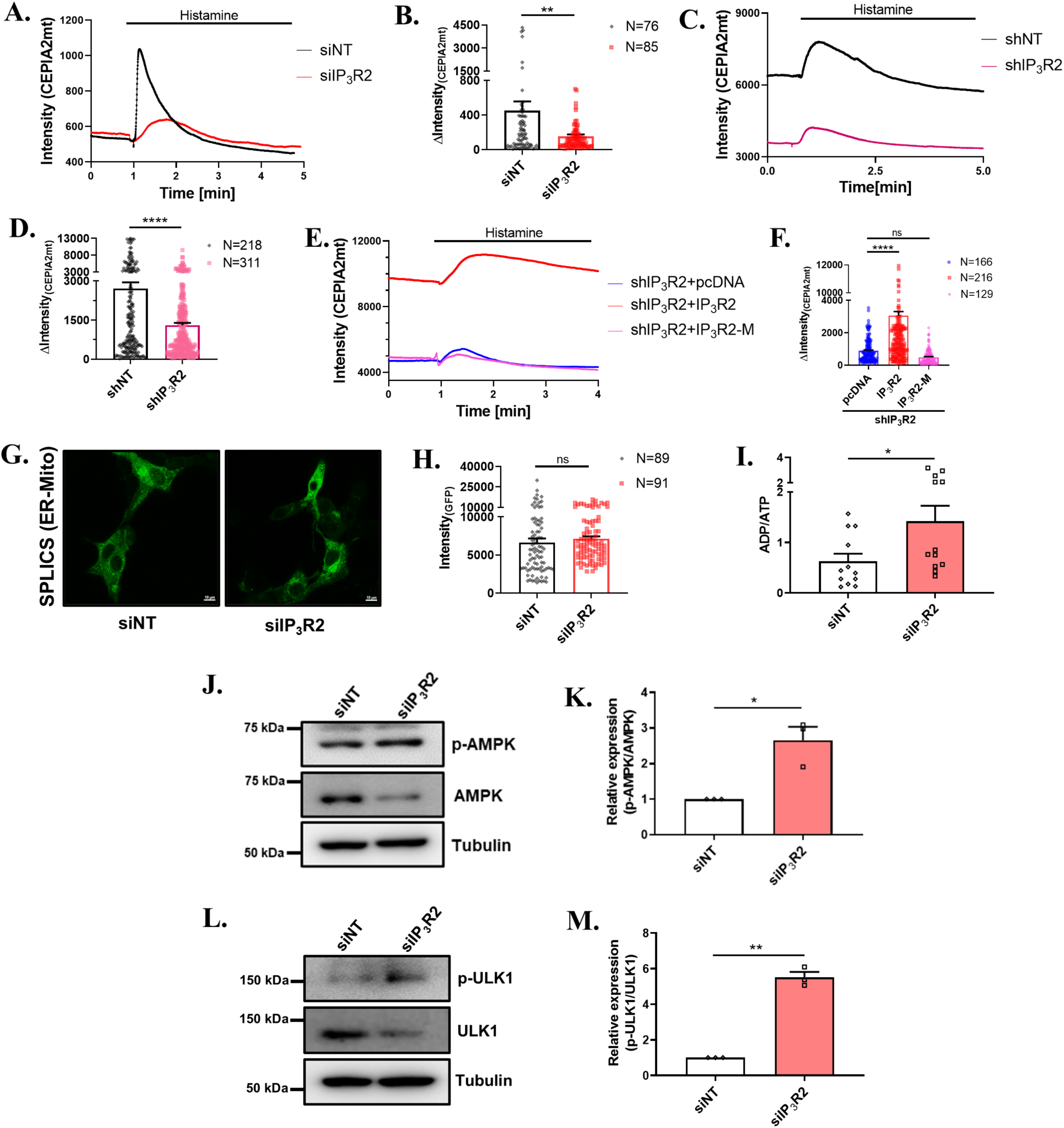
IP_3_R2 silencing reduces mitochondrial Ca^2+^ uptake and triggers melanophagy. (A) Representative mitochondrial Ca^2+^ imaging trace using the Cepia2mt probe in B16 cells stimulated with the Histamine after 72hrs of siRNA transfection. (B) Bar graph showing the quantification of Ca^2+^ imaging traces stimulated with 50µM histamine, where ‘N’ denotes the total number of regions of interest (ROI) in that trace. (C) Representative mitochondrial Ca^2+^ imaging trace using the Cepia2mt probe in B16 cells having stable IP_3_R2 knockdown upon stimulation with Histamine. (D) Bar graph showing the quantification of Ca^2+^ imaging traces stimulated with 50µM histamine, where ‘N’ denotes the total number of ROI in that trace. (E) Representative mitochondrial Ca^2+^ imaging trace using the Cepia2mt probe in B16 cells having stable IP_3_R2 knockdown demonstrating rescue with IP_3_R2 and IP_3_R2-M, upon stimulation with Histamine. (F) Bar graph showing the quantification of Ca^2+^ imaging traces stimulated with 50µM histamine, where ‘N’ denotes the total number of ROI in that trace. (G) Representative images of confocal imaging in B16 cells after transfected with siNT or siIP_3_R2 along with SPLICS Mt-ER Short P2A construct, scale bar, 10µm. (H) Bar graph showing the quantification of the GFP intensity, where ‘N’ denotes the number of cells imaged. (I) Bar graph shows quantification of the luminescent intensity measures changes in the ADP/ATP ratio (N=12). (J) Representative image of immunoblot showing expression of p-AMPK and AMPK in B16 cells after siRNA silencing of IP_3_R2. (K) Bar graph showing the densitometric analysis of the relative changes in the levels of p-AMPK to total AMPK (N=3). (L) Representative image of immunoblot showing expression of p-ULK1 and total ULK1 in B16 cells after siRNA silencing of IP_3_R2. (M) Bar graph showing the densitometric analysis of the relative changes in p-ULK1 to total ULK1 (N=3). Data presented are mean ± SEM. For statistical analysis, an unpaired t-test was performed for panels B, D, H, I, one sample t-test was performed for panels K and M and Dunnett’s multiple comparisons test was performed for panel F using GraphPad Prism software. Here, ‘ns’ means non-significant; * p <0.05, ** p < 0.01, and **** p < 0.0001.

We next examined whether the reduced mitochondrial Ca^2+^ uptake observed in IP_3_R2 knockdown cells is solely due to the loss of IP_3_R2 function or if it is accompanied with the changes in the ER and mitochondria proximity. We utilized a split-GFP-based fluorescent reporter system engineered to study ER and mitochondria proximity (Bartok *et al*, 2019; Cieri *et al*, 2017). We observed comparable fluorescence signals in the IP_3_R2 knockdown and control siNT conditions, suggesting no significant alteration in the ER-mitochondria contact sites upon IP_3_R2 silencing (**Fig 6G-H**). Taken together, our data highlights that the decrease in IP_3_R2 activity is the major contributor to the reduced mitochondrial Ca^2+^ uptake.

Literature suggests that decrease in mitochondrial Ca^2+^ uptake can impair mitochondrial function and diminish ATP production. This leads to an increase in the ADP/ATP or AMP/ATP ratio, which subsequently triggers activation of the autophagic pathway (Cárdenas *et al*, 2010; Sarkar *et al*, 2005; Antonia & Baldwin, 2018; Fink *et al*, 2017). Therefore, we assessed ADP/ATP ratio upon IP_3_R2 knockdown and observed a significant rise in the ADP/ATP ratio (**Fig 6I**). The increase in the ADP/ATP ratio is a hallmark of energetic stress and it amplifies phosphorylation of energy-sensing enzyme AMP-activated protein kinase (AMPK) (Viollet, 2017; Antonia & Baldwin, 2018). We examined the levels of phosphorylated AMPK and witnessed an elevated ratio of phosphorylated AMPK to total AMPK, suggesting that IP_3_R2 silencing enhances AMPK activity (**Fig 6J-K**). AMPK activation triggers macroautophagy through multiple signaling cascades to preserve cellular homeostasis and maintain energy balance. The key signaling modules working downstream of AMPK activation to drive macroautophagy are Unc-51-like kinase1 (ULK1) and mTOR pathway (Xu *et al*, 2024; Liu *et al*, 2024; Wang *et al*, 2022b). Therefore, we examined the status of these pathways upon IP_3_R2 silencing. Upon western blotting, we observed an increase in the ULK1 phosphorylation in IP_3_R2 silenced cells (**Fig 6L-M**). However, similar analysis revealed that IP_3_R2 silencing did not affect mTOR inactivation, as the phosphorylated mTOR/Total mTOR ratio was unaffected (**Supplementary Fig 4A-B**). Likewise, mTOR downstream targets, ribosomal protein S6 kinase beta-1 (p70S6K) and eukaryotic translation initiation factor 4E-binding protein 1 (4EBP1) remained unaffected (**Supplementary Fig 4C-F**). ULK1 phosphorylation induces autophagy by enabling the formation of autophagy initiation complex or ULK complex (**Fig 6L-M**) (Wang *et al*, 2018; Ryu *et al*, 2021). Taken together, our data suggests that IP_3_R2 silencing augments melanophagy, at least partially, by reducing mitochondrial Ca^2+^ uptake and thereby facilitating autophagy initiation complex formation.

### IP_3_R2 silencing decreases lysosomal pH

Since lysosomes play a crucial role in macroautophagy, we investigated potential impact of IP_3_R2 knockdown on lysosome biology. Lysosomal pH is a critical regulator of the autophagic cargo degradation as it controls activity of degradative enzymatic within lysosomes (Zeng *et al*, 2023). We estimated lysosomal pH using a ratiometric pH-sensitive fluorescent probe Lysosensor Yellow/Blue. We observed that IP_3_R2 silencing results in a significant decrease in lysosomal pH, as evident from elevated Yellow/Blue ratio (**Fig 7A-B**). To further validate this data, we utilized LAMP1-RpHLuorin2 ratiometric sensor to measure lysosomal pH. Our results show that IP_3_R2 silencing leads to a significant decrease in lysosomal pH, as indicated by a lower 405/488nm fluorescence ratio in the IP_3_R2 knockdown condition as compared to siNT control condition (**Fig 7C-D**). The data from two independent ratiometric probes demonstrates that IP_3_R2 silencing decreases lysosomal pH and that leads to enhanced melanophagy thereby reducing pigmentation. We further validated positive association between lysosomal pH and pigmentation levels by utilizing primary human melanocytes from two distinct origins i.e. Caucasian (lightly pigmented) and Afro-American (darkly pigmented). We observed that lightly pigmented primary human melanocytes exhibit a substantially lower lysosomal pH, as indicated by an increased Yellow/Blue ratio of Lysosensor Yellow/Blue, in comparison to darkly pigmented primary human melanocytes (**Supplementary Fig 5A-B**). Taken together, our data show that IP_3_R2 silencing reduces lysosomal pH, which is associated with lower pigmentation levels.

**Figure 7:**
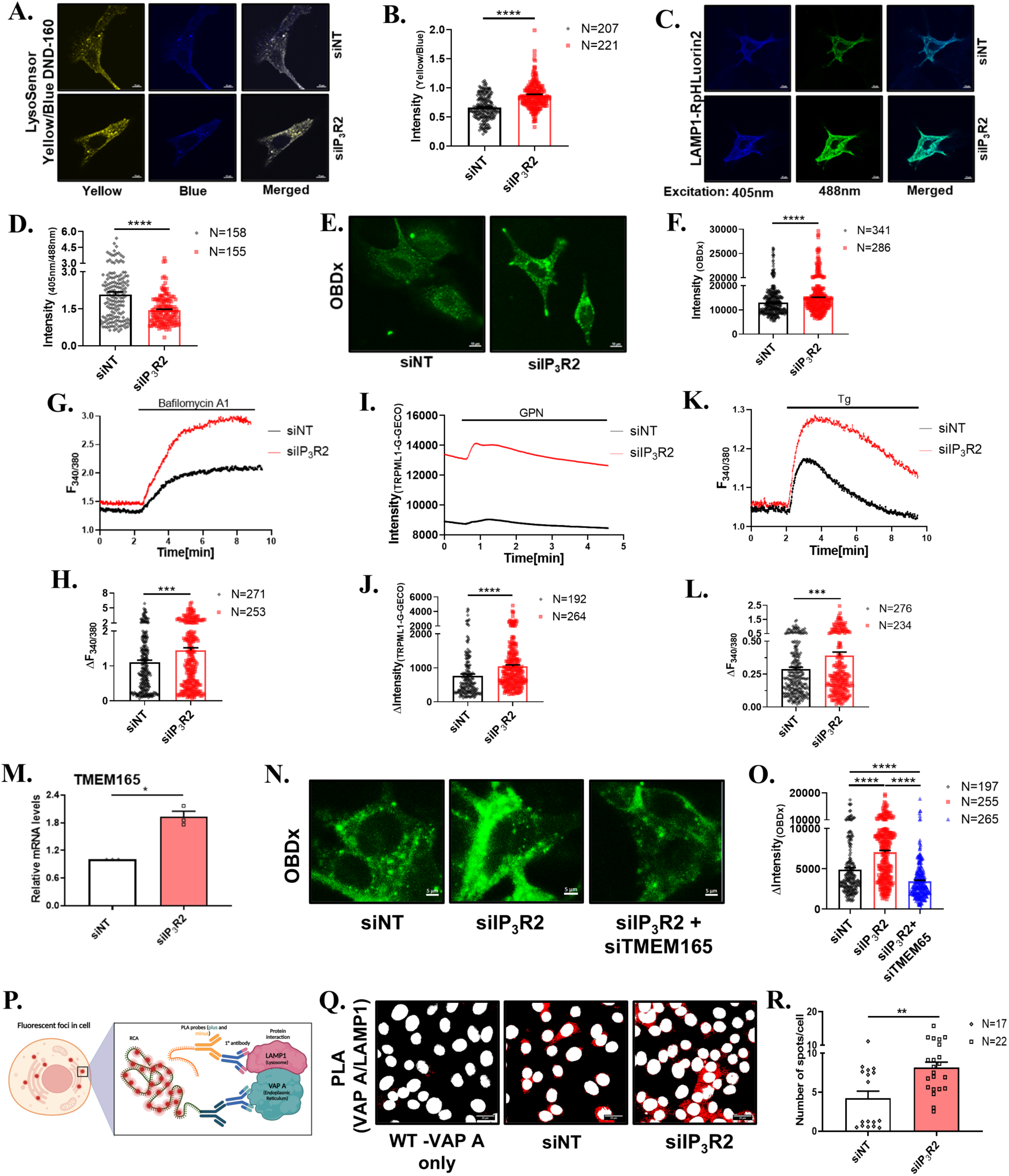
IP_3_R2 silencing decreases lysosomal pH and increases Ca^2+^ content in the lysosome. (A) Representative images of confocal imaging in B16 cells loaded with LysoSensor Yellow/Blue DND-160 dye for 5 mins after transfected with siNT or siIP_3_R2, scale bar, 10µm. (B) Bar graph showing the quantification of the intraluminal lysosomal pH in Yellow/Blue ratio, where ‘N’ denotes the number of cells imaged. (C) Representative images of confocal imaging in B16 cells after transfected with siNT or siIP_3_R2 along with LAMP1-RpHLuorin2 construct, scale bar, 10µm (D) Bar graph showing the quantification of the intraluminal lysosomal pH, in the 405nm:488nm ratio, which is alternatively excited at 405nm and 488nm, where ‘N’ denotes the number of cells. (E) Representative images of confocal imaging in B16 cells loaded with OG-BAPTA-dextran (OBDx) after transfected with siNT or siIP_3_R2, scale bar, 10µm. (F) Bar graph showing the quantification of the GFP intensity, where ‘N’ denotes the number of cells imaged. (G) Representative traces of Fura2AM based Ca^2+^ imaging in B16 cells stimulated with the BafilomycinA1 after 72hrs of siRNA transfection. (H) Bar graph showing the quantification of Ca^2+^ imaging traces stimulated with 1µM BafilomycinA1, where ‘N’ denotes the total number of cells imaged. (I) Representative Ca^2+^ imaging trace using the TRPML1-G-GECO probe in B16 cells stimulated with the GPN after 72hrs of siRNA transfection. (J) Bar graph showing the quantification of Ca^2+^ imaging traces stimulated with 300µM GPN, where ‘N’ denotes the total number of ROI in that trace. (K) Representative traces of Fura2AM based Ca^2+^ imaging in B16 cells stimulated with the Tg after 72hrs of siRNA transfection. (L) Bar graph showing the quantification of Ca^2+^ imaging traces stimulated with 2µM Tg, where ‘N’ denotes the total number of cells imaged. (M) qRT–PCR analysis showing increase in TMEM165 mRNA expression after 72hrs of siRNA mediated IP_3_R2 silencing. (N=3). (N) Representative images of confocal imaging in B16 cells loaded with OG-BAPTA-dextran (OBDx) after transfection with siRNA, scale bar, 5µm. (O) Bar graph showing the quantification of the GFP intensity, where ‘N’ denotes the number of cells imaged. (P) A schematic illustration demonstrating the mechanism of PLA by utilizing Primary antibody of LAMP1 and VAP A. (Q) Representative images of confocal imaging in B16 cells performing PLA between LAMP1 and VAP A proteins after transfected with siNT or siIP_3_R2, scale bar, 20µm. (R) Bar graph showing the quantification of the number of PLA spots per cell, where ‘N’ represents the number of images analyzed in the respective condition. Data presented are mean ± SEM. For statistical analysis, an unpaired t-test was performed for panels B, D, F, H, J, L and R, one sample t-test was performed for panel P, and Dunnett’s multiple comparisons test was performed for panel O using GraphPad Prism software. Here, * p <0.05, ** p < 0.01, *** p < 0.001 and **** p < 0.0001.

### IP_3_R2 silencing enhances lysosomal Ca^2+^ content

Given the crucial role of Ca^2+^ in modulating lysosomal function (Lloyd-Evans *et al*, 2010; Zou *et al*, 2015), we next examined the effect of IP_3_R2 knockdown on lysosomal Ca^2+^ levels. Earlier studies have shown that lysosomes accumulate Ca^2+^ released by all IP_3_ receptor subtypes, and blocking these receptors leads to lysosomal dysfunction and lysosomal storage disorders (LSD)-like phenotype (Garrity *et al*, 2016; Atakpa *et al*, 2018). Therefore, we expected that silencing the IP_3_R2 channel would reduce lysosomal Ca^2+^ levels. However, to our surprise, we observed that IP_3_R2 silencing increased Ca^2+^ content within the lysosomes in comparison to control condition, as measured with Oregon Green 488 BAPTA-1 dextran (OBDx) (**Fig 7E-F**). To further substantiate these findings, we used an indirect approach to evaluate lysosomal Ca^2+^ levels using Fura2-AM. We used an established protocol of releasing lysosomal Ca^2+^ into the cytosol by treating cells with bafilomycin A1. We observed that lysosomal Ca^2+^ release is higher in IP_3_R2 silenced cells compared to the siNT transfected control cells (**Fig 7G-H**). Moreover, we employed Glycyl-L-phenylalanine 2-naphthylamide (GPN) to disrupt the lysosomal membrane integrity and then measured the resultant Ca^2+^ elevation at the lysosomal periphery using the TRPML1-G-GECO Ca^2+^ sensor. This analysis also revealed a significantly greater Ca^2+^ release in the IP_3_R2 knockdown condition compared to the control (**Fig 7I-J**). Taken together, our live-cell Ca^2+^ imaging data from three independent experimental setups clearly demonstrate that IP_3_R2 silencing results in higher lysosomal Ca^2+^ accumulation. This further suggests that the higher lysosomal Ca^2+^ levels are associated with enhanced melanophagy and lower pigmentation levels. Consistent with these findings, we found that lightly pigmented primary human melanocytes have significantly elevated lysosomal Ca^2+^ content compared to darkly pigmented primary human melanocytes (**Supplementary Fig 5C-D**).

The above findings prompted an investigation into the mechanism by which lysosomes accumulate higher Ca^2+^ upon IP_3_R2 silencing. Recent studies have demonstrated that TMBIM6, a leaky channel on the ER membrane, facilitates Ca^2+^ uptake into lysosomes (Kim *et al*, 2021) and lysosomal transmembrane protein 165 (TMEM165) acts as a Ca^2+^ entry channel thereby bringing Ca^2+^ into lysosomes (Zajac *et al*, 2024). Therefore, we measured Ca^2+^ release from ER leaky channels using cytosolic Ca^2+^ indicator Fura2 AM. We treated B16 cells with thapsigargin, which inhibits the SERCA pump and we examined ER Ca^2+^ leak into cytosol. We observed an increase in ER Ca^2+^ leak in the IP_3_R2 knockdown cells compared to the non-targeting control (**Fig 7K-L**). Further, our qRT-PCR analysis revealed that TMEM165 expression is significantly elevated in IP_3_R2 silencing condition compared to the control (**Fig 7M**). To study the role of TMEM165 in IP_3_R2 silencing induced lysosomal Ca^2+^ influx, we silenced TMEM165 using siRNA. qRT-PCR analysis demonstrates decrease in TMEM165 expression upon TMEM165 silencing **(Supplementary Fig 6A)**. Thereafter, we measured lysosomal Ca^2+^ using OBDx and observed reduced Ca^2+^ in the lysosomal lumen upon TMEM165 silencing **(Supplementary Fig 6B-C)**. Importantly, along with reduced lysosomal Ca^2+^, TMEM165 knockdown resulted in an increase in pigmentation **(Supplementary Fig 6D-E)**. Next, we examined the role of TMEM165 in IP_3_R2 silencing induced increase in lysosomal Ca^2+^ levels by co-transfecting IP_3_R2 and TMEM165 siRNAs. We compared lysosomal Ca^2+^ content among control siNT, siIP_3_R2 only and siIP_3_R2+siTMEM165 conditions. This analysis demonstrated that IP_3_R2 knockdown driven rise in lysosomal Ca^2+^ is significantly reduced upon co-silencing of TMEM165 **(Fig 7N-O)**. These results suggest that the loss of IP_3_R2 leads to an increase in lysosomal Ca^2+^ uptake via lysosomal calcium importer channel TMEM165.

To investigate if proximity between ER and lysosomes influences Ca^2+^ transfer from ER to lysosome upon IP_3_R2 silencing, we employed in situ proximity ligation assays (PLA). This technique allows to determine if an ER protein, VAP-A and a lysosomal protein, LAMP1, are located within ∼40 nm of each other, indicating close spatial interactions (**Fig 7P**). The PLA data shows an increase in PLA spots in IP_3_R2 silenced condition compared to the control condition (**Fig 7Q-R**), suggesting that IP_3_R2 downregulation increases ER and lysosomal interactions.

Taken together, our data demonstrate that IP_3_R2 silencing leads to elevated Ca^2+^ levels within ER. This subsequently enhances Ca^2+^ release through leaky ER channels and increases Ca^2+^ influx into lysosomes via TMEM165. Moreover, our data show that IP_3_R2 knockdown augments ER and lysosomal interaction thereby contributing to increased lysosomal Ca^2+^ levels. Collectively, these results reveal that the loss of IP_3_R2 results in a substantial increase in lysosomal Ca^2+^ level, which may serve as a critical upstream signal that triggers melanophagy.

### Interplay between IP_3_R2 and TRPML1 drives lysosomal Ca^2+^ release and melanophagy

Recent studies have reported that lysosomal Ca^2+^ efflux via TRPML channels triggers autophagy (Medina *et al*, 2015; Scotto Rosato *et al*, 2019). Moreover, literature suggests that the acidic environment within the lysosome stimulates TRPML1 activity (Miao *et al*, 2015; Xiong & Zhu, 2016; Li *et al*, 2017). Since we observed an increase in lysosomal Ca^2+^ levels and lower lysosomal pH upon IP_3_R2 silencing, we investigated role of TRPML channels in driving melanophagy. Our FURA2-AM based cytosolic Ca^2+^ measurements revealed that stimulation with TRPML agonist MLSA1 results in higher lysosomal Ca^2+^ release in IP_3_R2 knockdown condition compared to siNT control (**Fig 8A-B**). We next corroborated role of TRPML1 in enhanced lysosomal Ca^2+^ release by using a genetically encoded Ca^2+^ sensor TRPML1-G-GECO that measures Ca^2+^ levels in TRPML1 proximity (Davis *et al*, 2020). The live-cell Ca^2+^ imaging with TRPML1-G-GECO demonstrated that MLSA1 stimulation results in higher Ca^2+^ release from TRPML1 in IP_3_R2 knockdown condition (**Fig 8C-D**). This data suggests that IP_3_R2 downregulation augments TRPML1 activity.

**Figure 8:**
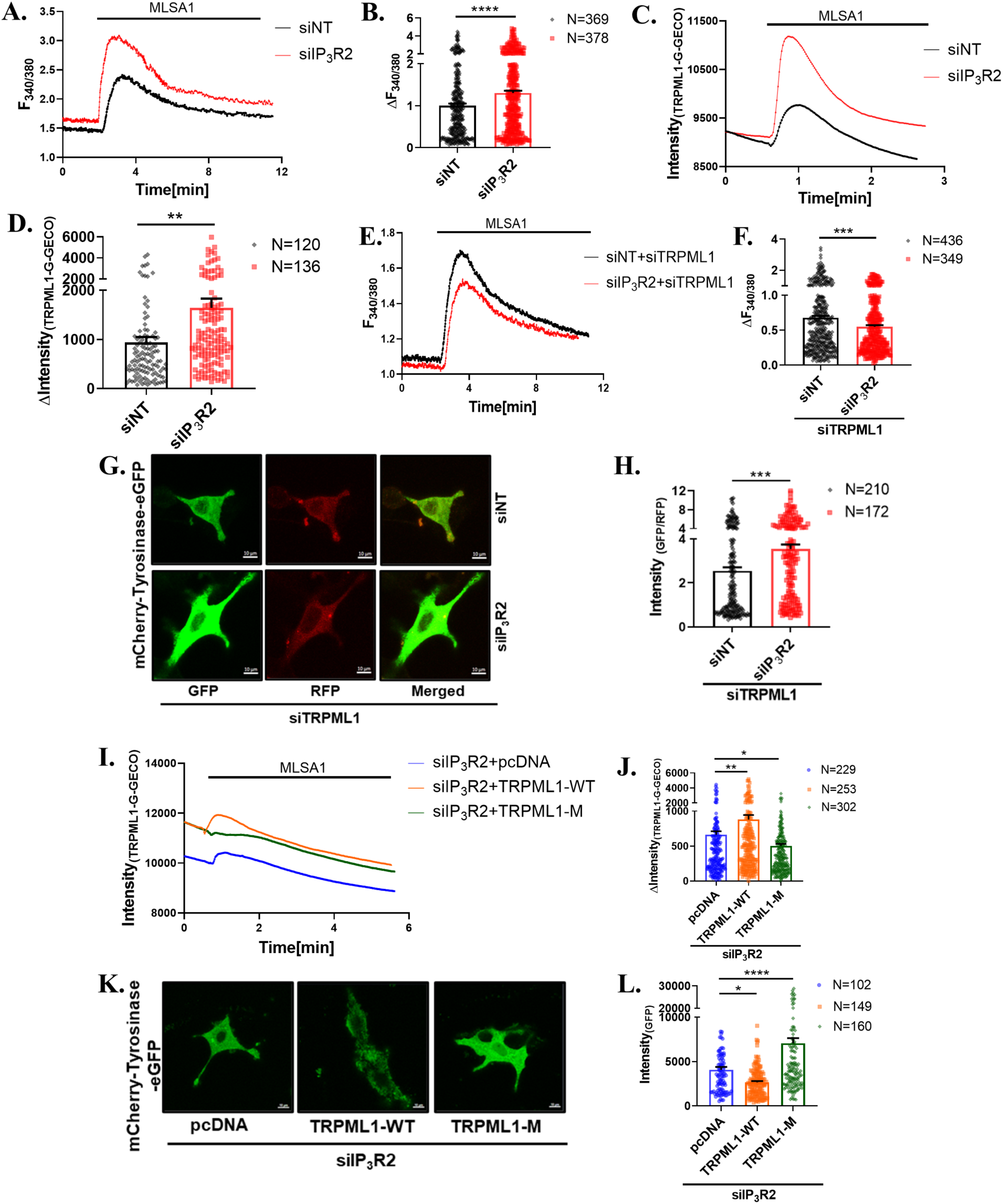
IP_3_R2 knockdown augments TRPML1 activity and enhances melanophagy flux. (A) Representative traces of Fura2AM based Ca^2+^ imaging in B16 cells stimulated with the MLSA1 after 72hrs of siRNA transfection. (B) Bar graph showing the quantification of Ca^2+^ imaging traces stimulated with 20µM MLSA1, where ‘N’ denotes the total number of cells in that trace. (C) Representative Ca^2+^ imaging trace using the TRPML1-G-GECO probe in B16 cells stimulated with the MLSA1 after 72hrs of siRNA transfection. (D) Bar graph showing the quantification of Ca^2+^ imaging traces stimulated with 20µM MLSA1, where ‘N’ denotes the total number of ROI in that trace. (E) Representative traces of Fura2AM based Ca^2+^ imaging in B16 cells stimulated with the MLSA1 after 72hrs of siRNA mediated dual transfections of IP_3_R2 and TRPML1. (F) Bar graph showing the quantification of Ca^2+^ imaging traces stimulated with 20µM MLSA1, where ‘N’ denotes the total number of cells in that trace. (G) Representative images of confocal imaging in B16 cells after co-transfected with siIP_3_R2 and siTRPML1 along with mCherry-Tyrosinase-EGFP probe, scale bar, 10µm. (H) Bar graph shows the quantification of the autolysosome intensity in the GFP:RFP ratio, where ‘N’ denotes the number of cells. (I) Representative trace of lysosomal Ca^2+^ imaging using the TRPML1-G-GECO probe in B16 cells transfected with siIP_3_R2 demonstrating rescue with TRPML1-WT and TRPML1-M. (J) Bar graph showing the quantification of Ca^2+^ imaging traces stimulated with 20µM MLSA1, where ‘N’ denotes the total number of ROI in that trace. (K) Representative images of confocal imaging using the mCherry-Tyrosinase-EGFP construct in B16 cells transfected with siIP_3_R2 demonstrating rescue with TRPML1-WT and TRPML1-M, scale bar, 10µm. (L) Bar graph showing the quantification of the GFP intensity (mCherry inherently present in the TRPML1 plasmid), where ‘N’ denotes the number of cells imaged. Data presented are mean ± SEM. For statistical analysis, unpaired t-test was performed for panels B, D, F, H, and Dunnett’s multiple comparisons test was performed for panels J, L using GraphPad Prism software. Here, * p <0.05, ** p < 0.01, *** p < 0.001 and **** p < 0.0001.

We then carried out TRPML1 loss of function studies. We initially conducted qRT-PCR analysis to evaluate the specificity of siTRPML1 across different TRPML isoforms. This analysis demonstrated that siRNA mediated TRPML1 knockdown is both robust and specific (**Supplementary Fig 7A**). Using the TRPML1-G-GECO Ca^2+^ sensor, we measured MLSA1 stimulated Ca^2+^ release from TRPML1 and observed almost complete abrogation of TRPML1 mediated lysosomal Ca^2+^ release in siTRPML1 condition (**Supplementary Fig 7B-C**). This data demonstrates that the siTRPML1 efficiently and specifically decreases TRPML1 expression and activity. We next validated TRPML1’s role in enhanced lysosomal Ca^2+^ release upon IP_3_R2 knockdown by co-silencing IP_3_R2 and TRPML1. We then assessed lysosomal Ca^2+^ release from these cells and the control (co-transfected with siNT and siTRPML1) cells. Our results show that the combined IP_3_R2 and TRPML1 knockdown leads to a decrease in MLSA1 stimulated lysosomal Ca^2+^ release (**Fig 8E-F**), in contrast to the increased Ca^2+^ release observed with IP_3_R2 silencing alone (**Fig 8A-D**). Taken together, our data reveals that IP_3_R2 knockdown enhances TRPML1 mediated lysosomal Ca^2+^ release.

We next examined role of TRPML1 in regulating lysosomal pH. Our analysis using LysoSensor Yellow/Blue revealed that TRPML1 knockdown decreases the Yellow/Blue ratio, suggesting an increase in lysosomal pH (**Supplementary Fig 7D-E**). Furthermore, we assessed TRPML1’s role in melanophagy using mCherry-Tyrosinase-eGFP probe. TRPML1 knockdown did not change melanophagy flux compared to siNT control (**Supplementary Fig 7F-G**). Therefore, our data suggests that TRPML1 knockdown alone does not impact melanophagy. We then investigated if TRPML1 contributes to the increase in melanophagy observed upon IP_3_R2 knockdown. We co-silenced IP_3_R2 and TRPML1 and assessed melanophagy with mCherry-Tyrosinase-eGFP probe. Our results show that the combined IP_3_R2 and TRPML1 knockdown leads to a reduction in melanophagy in comparison to the control condition (**Fig 8G-H**). To further validate role of TRPML1 in IP_3_R2 silencing induced melanophagy, we performed rescue experiments with either wild-type TRPML1 or TRPML1 non-conducting pore mutant (D471K/D472K) that cannot facilitate Ca^2+^ release from lysosomes (Pryor *et al*, 2006). Firstly, we corroborated the functionality of wild-type TRPML1 and pore-dead TRPML1 mutant (TRPML1-M) by measuring TRPML1 mediated lysosomal Ca^2+^ release with TRPML1-G-GECO. Our results show that overexpression of the wild-type TRPML1 leads to an increase in Ca^2+^ release compared to the control condition, while the pore-dead TRPML1 mutant exhibited a decrease in Ca^2+^ release relative to the control (**Supplementary Fig 7H-I**). Next, we examined if TRPML1 contributes to enhanced lysosomal Ca^2+^ release observed upon IP_3_R2 knockdown. We ectopically expressed either wild-type TRPML1 or TRPML1-M in IP_3_R2 silenced cells and measured Ca^2+^ release using TRPML1-G-GECO. Our data show that wild-type TRPML1 augments lysosomal Ca^2+^ release while TRPML1-M decreases the Ca^2+^ release in comparison to vector control pcDNA in IP_3_R2 silenced cells (**Fig 8I-J**). This data corroborates that TRPML1 mediated Ca^2+^ release contributes to enhanced lysosomal Ca^2+^ release observed upon IP_3_R2 silencing. We then investigated role of TRPML1 in driving melanophagy upon IP_3_R2 knockdown. We assessed melanophagy levels using the mCherry-Tyrosinase-eGFP probe. Our confocal analysis demonstrate that in IP_3_R2 silenced cells, overexpression of wild-type TRPML1 further enhanced melanophagy while ectopic expression of TRPML1-M led to a significant decrease in melanophagy (**Fig 8K-L**). These results clearly establish that downstream of IP_3_R2 knockdown, TRPML1 mediated lysosomal Ca^2+^ release plays a critical role in driving melanophagy.

### TRPML1 mediated lysosomal Ca^2+^ release regulates melanophagy via TFEB translocation

Our data clearly demonstrates that downstream of IP_3_R2 silencing TRPML1 facilitates augmented lysosomal Ca^2+^ release and enhanced melanophagy. Next, we investigated the molecular mechanism through which TRPML1 drives melanophagy. Recent literature suggests that TRPML1 mediated lysosomal Ca^2+^ release activates autophagy via transcription factor EB (TFEB) nuclear translocation (Park *et al*, 2022; Medina *et al*, 2015; Scotto Rosato *et al*, 2019; Roczniak-Ferguson *et al*, 2012). Therefore, we investigated the potential role of TFEB in enhancing melanophagy upon IP_3_R2 knockdown. Our confocal microscopy data using pEGFP-N1-TFEB shows an increase in TFEB nuclear localization in IP_3_R2 silenced condition compared to the control condition (**Fig 9A-B**). We further validated this observation by performing subcellular fractionation followed by probing nuclear to cytosolic ratio of TFEB via immunoblotting. This analysis revealed an increase in nuclear to cytoplasmic ratio of TFEB in the IP_3_R2 knockdown condition compared to control (**Fig 9C-D**). Collectively, the microscopy and biochemical findings show that there is elevated TFEB nuclear translocation upon IP_3_R2 silencing suggesting that it can be a likely mechanism underlying the enhanced melanophagy observed upon IP_3_R2 downregulation.

**Figure 9:**
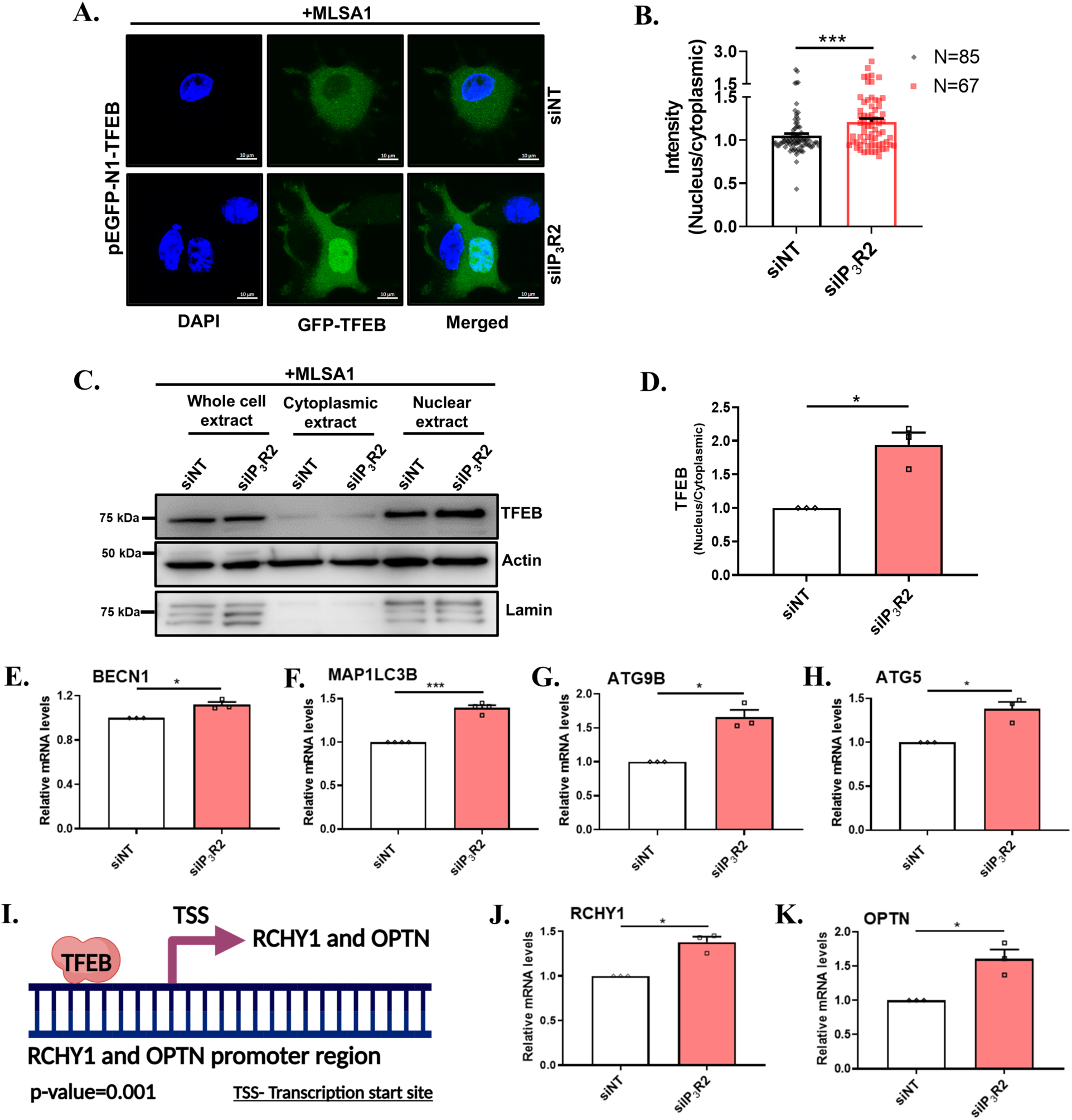
TRPML1 mediated lysosomal Ca^2+^ release regulates melanophagy via TFEB nuclear translocation. (A) Representative images of confocal imaging in B16 cells demonstrates TFEB nuclear translocation stimulated with 20µM MLSA1 for 1 hour, after transfecting with siNT or siIP_3_R2 along with pEGFP-N1-TFEB construct, scale bar, 10µm. (B) Bar graph showing the quantification of the GFP intensity from the cytoplasm and nucleus in the nucleus/cytoplasmic ratio following the addition of 20µM MLSA1 for 1hr, where ‘N’ denotes the number of cells imaged. (C) Representative image of immunoblot showing expression of TFEB in B16 cells from the cytoplasm and nucleus following subcellular fractionation after siRNA silencing of IP_3_R2. (D) Bar graph showing the densitometric analysis of the relative changes in the levels of TFEB by quantitating nucleus/cytoplasmic ratio (N=3). (E) qRT PCR analysis showing increase in BECN1 mRNA expression after 72hrs of siRNA mediated IP_3_R2 silencing. (N=3). (F) qRT PCR analysis showing increase in MAP1LC3B mRNA expression after 72hrs of siRNA mediated IP_3_R2 silencing. (N=4). (G) qRT PCR analysis showing increase in ATG9B mRNA expression after 72hrs of siRNA mediated IP_3_R2 silencing. (N=3). (H) qRT PCR analysis showing increase in ATG5 mRNA expression after 72hrs of siRNA mediated IP_3_R2 silencing. (N=3). (I) A schematic illustration demonstrating the transcriptional regulation of the TFEB on expression of genes RCHY1 and OPTN. (J) qRT PCR analysis showing increase in RCHY1 mRNA expression after 72hrs of siRNA mediated IP_3_R2 silencing. (N=3). (K) qRT PCR analysis showing increase in OPTN mRNA expression after 72hrs of siRNA mediated IP_3_R2 silencing. (N=3). Data presented are mean ± SEM. For statistical analysis, one sample t-test was performed for panels B, D, E, F, G, H, J, K using GraphPad Prism software. Here, * p <0.05, *** p < 0.001.

Previous studies have demonstrated that TFEB enhances transcription of several autophagic genes (Medina *et al*, 2015; Palmieri *et al*, 2017; Di Malta *et al*, 2019). We performed qRT-PCR analysis in IP_3_R2 silenced and corresponding control cells. We observed that IP_3_R2 silencing led to an increase in the mRNA levels of several genes (Beclin1, MAP1LC3B, ATG9B and ATG5) involved in the initiation and progression of autophagy (**Fig 9E-H**). Recently, ring finger and CHY Zinc finger domain containing 1 (RCHY1) and optineurin (OPTN) were reported as key regulators of melanophagy (Lee *et al*, 2024). We next examined if TFEB transcriptionally controls RCHY1 and OPTN to drive melanophagy. First, we performed extensive bioinformatic analysis using the Eukaryotic Promoter Database at a very stringent p-value threshold of 0.001 and found that TFEB has multiple potential binding sites on RCHY1 and OPTN promoters (**Fig 9I**). We corroborated this observation via qRT-PCR experiments, which revealed an increase in RCHY1 and OPTN expression upon IP3R2 knockdown suggesting that TFEB-mediated upregulation of these genes is involved in the melanophagy (**Fig 9J-K**). Collectively, our data reveals that upon IP_3_R2 knockdown there is enhanced TRPML1 mediated lysosomal Ca^2+^ release, which leads to TFEB nuclear translocation and transcriptional upregulation of autophagy and melanophagy regulators.

In summary, our work reveals that IP_3_R2 acts a critical driver of pigmentation by keeping a check on melanophagy. Our *in vitro* studies in multiple independent pigmentation models and *in vivo* experiments in zebrafish demonstrate that IP_3_R2’s ER Ca^2+^ release function is required for controlling pigmentation levels. Further, we report that IP_3_R2 mediated ER-mitochondrial and ER-lysosomal crosstalk is a crucial regulator of melanophagy. Our data show that IP_3_R2 silencing decreases mitochondrial Ca^2+^ whereas it increases lysosomal Ca^2+^ levels. This in turn induces melanophagy via AMPK-ULK and TRPML1-TFEB signaling cascades respectively. Hence, this work uncovers that IP_3_R2-mediated Ca^2+^ signaling across organelles is a critical determinant of melanophagy and consequently skin pigmentation. Therefore, this signaling module offers potential therapeutic options for the management of pigmentary disorders and skin malignancies.

## Discussion

Inter-organelle communication is emerging as a key determinant of cellular physiology and a critical regulator of pathological outcomes. However, role of inter-organelle crosstalk in melanocyte function and pigmentation biology remains poorly understood. Disruptions in pigmentation pathways contribute to the development of pigmentary disorders, such as vitiligo and melasma. Further, they predispose to skin cancers, which are highly metastatic and associated with poor prognosis. Pigmentation is an outcome of balance between melanosome biogenesis and degradation. Although signaling modules that drive melanosome biogenesis are partially understood, the molecular mechanisms regulating melanophagy process remain largely unappreciated. Here, we reveal that IP_3_R2 positively regulates pigmentation by controlling melanophagy (**Fig. 10**). IP_3_R2 mediated ER-mitochondrial and ER-lysosomal Ca^2+^ signaling is a crucial modulator of melanosome degradation. IP_3_R2 silencing decreases mitochondrial Ca^2+^ uptake (**Fig. 6**) while it increases lysosomal Ca^2+^ concentration (**Fig. 7**). This in turn collectively enhances melanophagy thereby resulting a decrease in pigmentation.

**Figure 10:**
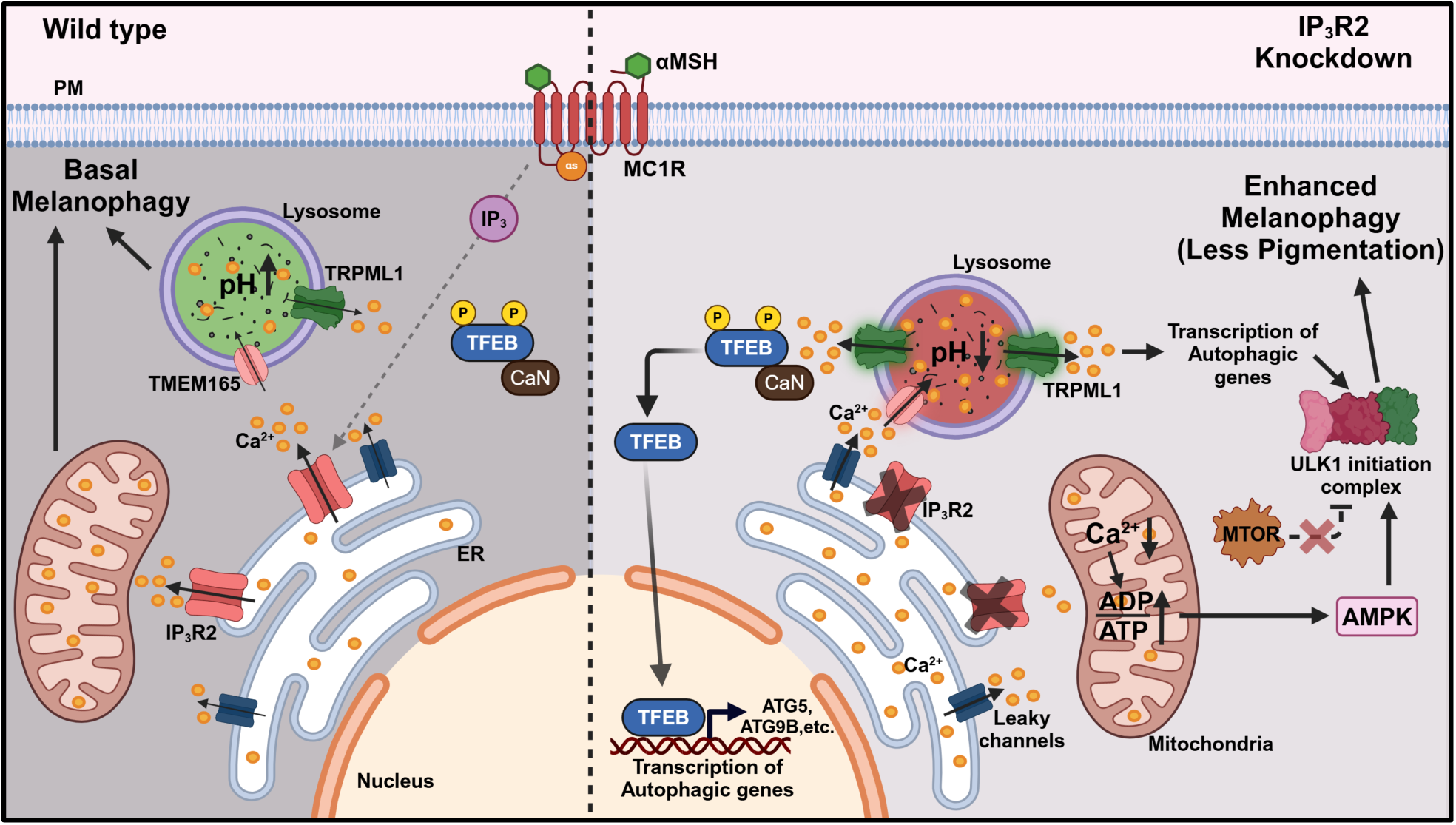
Multifaceted regulation of melanophagy mediated by IP_3_R2 knockdown.

Our data demonstrate that IP_3_R2 knockdown leads to an increase in stability of melanogenic proteins despite a concurrent decrease in pigmentation (**Fig. 3 and 4**). We performed live cell melanophagy flux assays using two *de novo* generated ratiometric probes to show that IP_3_R2 silencing induces melanophagy (**Fig. 5**). We further validated these results by performing biochemical assays (**Fig. 5**) and ultrastructural studies (**Supplementary Fig. 3)**. Mechanistically, the IP_3_R2 knockdown stimulates melanophagy via AMPK-ULK and TRPML1-TFEB signaling cascades that work downstream of ER-mitochondria and ER-lysosomal Ca^2+^ signaling respectively. Interestingly, the role of ER Ca^2+^ transients was recently reported in stimulating autophagosome formation (Zheng *et al*, 2022, 2025). It was showed that IP3R2 silencing in Cos7 kidney cells increases lysosomal Ca^2+^ stores. This in turn led to enhanced lysosomal Ca^2+^ efflux that initiates formation of FIP200 puncta on ER membrane. These ER associated FIP200 are crucial for autophagosome formation (Zheng *et al*, 2022). Although it was suggested that ER Ca^2+^ transients regulate lysosomal Ca^2+^ content and lysosomal Ca^2+^ efflux, the underlying molecular mechanisms responsible for increase in lysosomal Ca^2+^ stores and augmented lysosomal Ca^2+^ release remain unappreciated. Here, we show that IP_3_R2 silencing bring ER-lysosome closer and that results in higher lysosomal Ca^2+^ uptake via TMEM165 (**Fig. 7**). Further, we demonstrate IP_3_R2 knockdown results in a decrease in lysosomal pH, which subsequently enhances the activity of the TRPML1 channel resulting in higher lysosomal Ca^2+^ release (**Fig. 7**). Importantly, the co-silencing of IP_3_R2 and TRPML1 reduced lysosomal Ca^2+^ efflux measured via TRPML1-G-GECO indicating that TRPML1 plays a crucial role in mediating the Ca^2+^ release from lysosomes downstream of IP_3_R2 downregulation (**Fig. 7**). Further, TRPML1 mediated lysosomal Ca^2+^ release drives melanophagy (**Fig. 8**) by enhancing TFEB nuclear translocation and transcription of autophagy/melanophagy genes (**Fig. 9**).

Although lysosomes have other Ca^2+^ release channels such as TPCs and P2X4 channels, they most likely do not contribute to melanophagy. Indeed, literature suggest that TPCs do not contribute to TFEB translocation and thereby they would most likely not regulate melanophagy (Medina *et al*, 2015). Likewise P2X4 and TRPML2/3 channels get activated at high pH and are inactivated at low pH (Huang *et al*, 2014; Grimm *et al*, 2012) whereas IP_3_R2 silencing decreases lysosomal pH. It is important to highlight that TRPML1 is activated at low pH (Schmiege *et al*, 2017; Shen *et al*, 2012) and therefore, it is stimulated downstream of IP_3_R2 knockdown.

Interestingly, an earlier study reported that TRPML3 gain of function mutant mice show depigmentation of fur and tail (Xu *et al*, 2007). Though authors suggested loss of melanocyte function as a possible driver of depigmentation, the underlying molecular mechanism that induce depigmentation in this mice remain elusive. In future, it would be interesting to study the role of TRPML3 mediated lysosomal Ca^2+^ efflux in melanophagy to examine if it leads to depigmentation by enhancing melanophagy. Secondly, in the future studies, it would be worth to generate a TRPML1 gain of function mice and investigate its effect on mice pigmentation.

Taken together, this study reveals an intricate interplay between IP_3_R2-mediated Ca^2+^ dynamics across organelles and melanophagy. Our work show that IP_3_R2 silencing reduces mitochondrial Ca^2+^ uptake and simultaneously increases lysosomal Ca^2+^ levels. This in turn enhances melanophagy via AMPK-ULK and TRPML1-TFEB signaling modules respectively. In summary, we demonstrate that Ca^2+^ signaling across organelles is a crucial regulator of melanophagy and thereby pigmentation. Hence, this study holds the potential to unlock new avenues for treating pigmentary disorders and developing targeted strategies to manage skin malignancies.

## Materials and Methods

### Cell culture

B16-F10 murine melanoma cells were procured from the American Type Culture Collection and cultured in Dulbecco’s Modified Eagle’s Medium-high glucose (DMEM-HG; Sigma-Aldrich, D5648) supplemented with 10% fetal bovine serum (FBS; Gibco, 10270106), 1X Antibiotic Antimycotic (Thermo Fisher Scientific, 15240062). The B16-F10 cells were maintained at 60-80% confluency in a humidified incubator with 5% CO2 levels. Additionally, Human Epidermal Melanocytes, neonatal, lightly pigmented donor (HEMn-LP; Gibco, C0025C) were cultured in Medium 254 (M254; Gibco, M254CF) supplemented with human melanocyte growth supplement-2 (HMGS-2; Gibco, S0165) 1X Antibiotic Antimycotic (Thermo Fisher Scientific, 15240062) under the same environmental conditions. Experiments were conducted using cells between passages 3 to 6. Essential cell culture reagents used during experiments were phosphate-buffered saline, pH 7.2 (PBS; HIMEDIA, M1452), Trypsin (2.5%), and no phenol red (Gibco, 15090046).

### Low-density (LD) pigmentation model system

B16 cells were seeded at an initial density of 100 cells per/cm^2^ in Dulbecco’s Modified Eagle’s Medium supplemented with 10% fetal bovine serum, as described previously (Natarajan *et al*, 2014b; Motiani *et al*, 2018). The cells were then permitted to develop pigmentation gradually, and the experiments were terminated on Day 6 or Day 7.

### siRNA Transfection

Murine B16 melanoma cells were seeded at a density of 60,000 cells per well in a 6-well culture plate. The next day, transfection was performed using 50nM of siRNA (see Table 1) incubated with TurboFect Transfection Reagent (Thermo Fisher Scientific, R0531) for 30 min in 1:3 ratio (v/v) in Opti-MEM Reduced Serum Medium (Gibco,31985070). The transfection mixture was then added to the wells of 6 well plates. After 24 hours, melanocyte-stimulating hormone (αMSH; Sigma Aldrich, M4135) was added for 48 hours, followed by cell harvesting. If not otherwise stated.

**Table 1:**
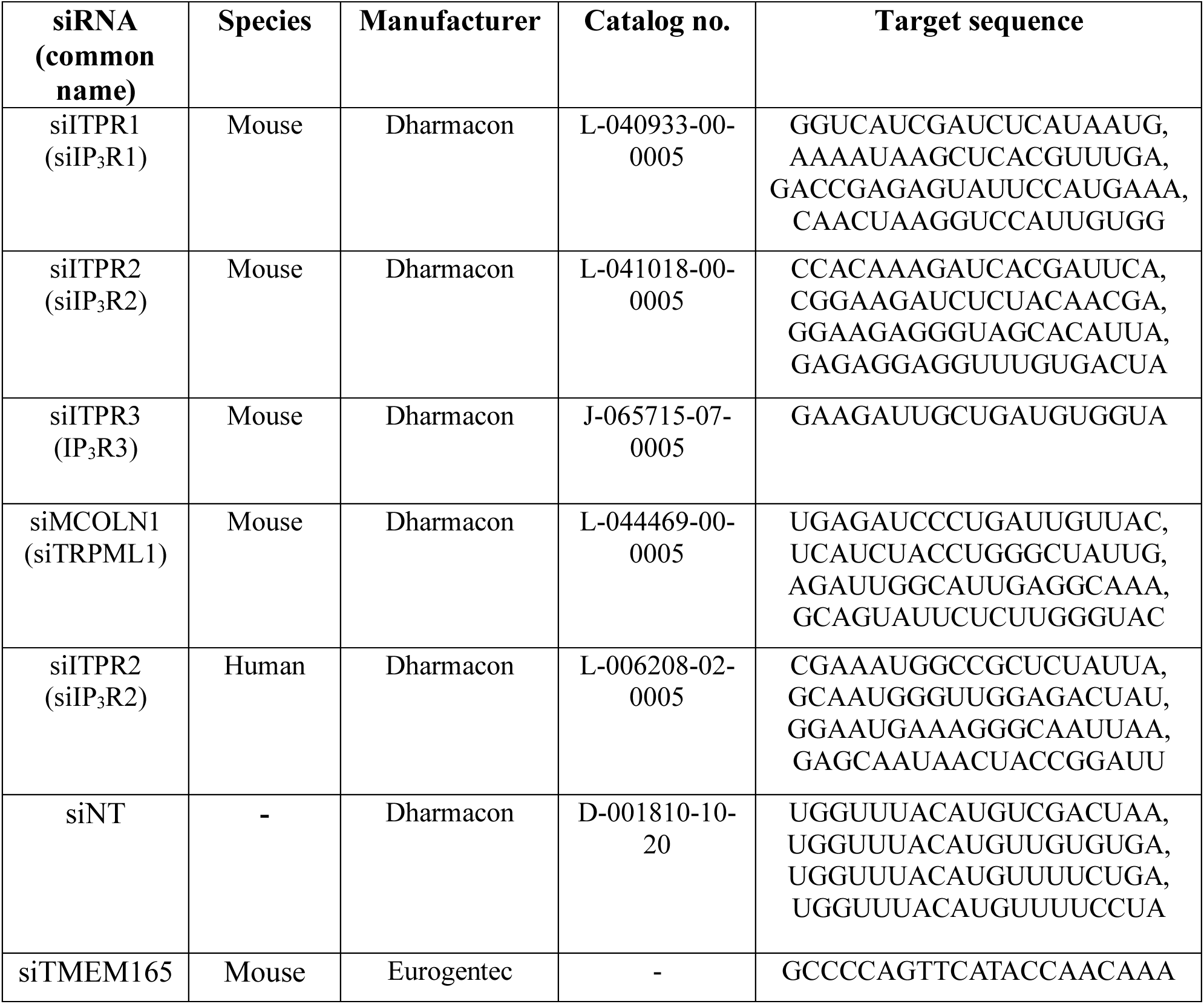
List of siRNA.

On day 3 of the low-density pigmentation model, 100 nM of siRNA (see Table 1) was incubated with DharmaFECT 2 Transfection Reagent (Dharmacon, T-2002-03) at a 1:3 ratio (v/v) in Opti-MEM Reduced Serum Medium for 45 minutes. The original LD day 3 media was saved, and the transfection mixture was added to the cells for 6 hours. Subsequently, the saved day 3 media was replaced, with the addition of 1µM αMSH. The cells were harvested on Day 6.

For HEMn-LP, the cells were first trypsinized, and a cell pellet containing 7-10 x 10^5 cells was obtained. The cells were then resuspended in 100uL of the Nucleofector Solution for Human Melanocytes – Neonatal, provided in the Human Melanocyte - Neonatal Nucleofector Kit. (Lonza, VPD1003). Subsequently, 5uM siRNA (see Table 1) was added to this cell suspension. The cell-siRNA mixture was transferred to an Aluminum Cuvette (included in the kit) and nucleofection was performed using the U-024 program of the Nucleofector™ 2b device. After the electroporation, the cells were resuspended in Medium 254 supplemented with Human Melanocyte Growth Supplement-2 and then seeded in a culture flask for a 72-hour incubation period.

### Lentiviral Stable Cell Line Generation

To achieve stable knockdown, mouse-specific short hairpin RNAs (shRNAs) targeting the non-targeting control (shNT) and IP_3_R2 (shITPR2) were obtained from Transomics Technologies and cloned into the lentiviral vector pGIPZ-mCMV-mCherry-Puromycin. As previously reported, the lentiviral constructs pCMV-VSVG (Addgene, 8454), pCMV-dR8.2 (Addgene, 8454), and pGIPZ-shNT/shITPR2 (Transomics Technologies, TLMSU1452) were mixed in a 1:3:2 ratio and combined with Lipofectamine 2000 (Thermo Fisher Scientific, 11668019) at a 1:2 ratio (w/v) in Opti-MEM medium (Arora *et al*, 2021; Tanwar *et al*, 2022a). The mixture was incubated for 45 minutes and then added to a flask containing 90% confluent HEK293T cells for 6 hours. The transfection mixture was then removed, and DMEM-HG supplemented with 2% Tet-negative fetal bovine serum was added. Viral particles were collected 48 hours post-transfection and concentrated using Amicon filters through centrifugation at 4000 rpm for 30 minutes in a swinging bucket. The concentrated viral particles, along with 10 µg/mL polybrene, were then added to B16 cells seeded at 50% confluency in a T25 flask for a 48-hour transduction period. Subsequently, 5 µg/mL of puromycin was applied to select the transduced cells. The knockdown was validated through Western blot analysis.

### RNA extraction and Real-time PCR

Total RNA was isolated from the cell samples using the RNeasy Mini Kit (Qiagen, 74104) following the manufacturer’s protocol. Complementary DNA was then synthesized from the extracted RNA through reverse transcription using the High-Capacity cDNA Reverse Transcription Kit (Thermo Fisher Scientific, 4368814) following the manufacturer’s protocol. Subsequently, quantitative real-time PCR with complimentary DNA (cDNA) was conducted using SYBR green (Takara, RR420A), following the manufacturer’s protocol utilizing the Quant Studio 6 Flex from Applied Biosystems or CFX96 Touch Real-Time PCR Detection System from Bio-Rad. The expression analysis was normalized against the GAPDH housekeeping gene unless otherwise stated. Primers used in this study are listed in Table 2.

**Table 2:**
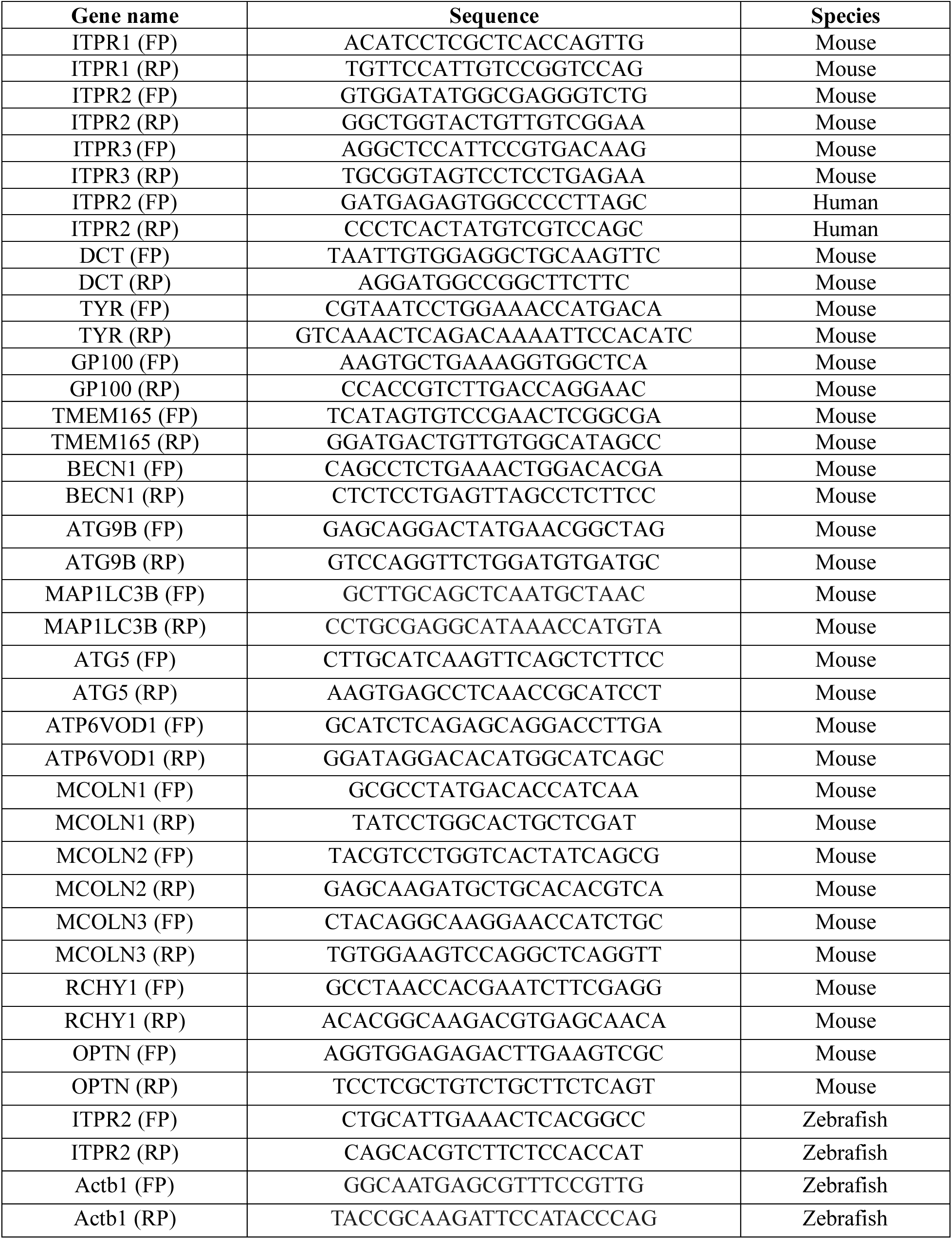
List of qRT-PCR primers (FP stands for forward and RP for reverse primer).

### Antibodies and Western blot

Total cellular protein extracts were prepared by solubilizing the samples in a buffer containing Membrane-bound Protein Extraction Buffer (10mM Tris-HCl, 100mM NaCl, 1mM EDTA, 1mM EGTA, 1mM NaF, 20mM Na4P2O7, 2mM Na3VO4, 1% Triton X-100 (v/v), 10% Glycerol, 0.5% sodium deoxycholate, 0.01% SDS (w/v), at pH 7.4) and cOmplete Mini, EDTA-free Protease Inhibitor Cocktail (Roche, 4693159001) with incubation for 30 minutes at 4°C. The soluble fraction was then recovered in the supernatant after spinning at 15,000 rpm for 15 minutes at 4°C, and the total protein concentration was quantified using the Pierce BCA Protein Assay Kits (Thermo Fisher Scientific, 23227).

Proteins were resolved using 8-14% gel using the SDS-PAGE method, then transferred to a 0.45µ Polyvinylidene fluoride (PVDF) membrane using the Mini-PROTEAN Tetra electrophoresis system (BioRad). The PVDF membrane was blocked in Tris-buffered saline containing 0.1% Tween-20 and 5% BSA or non-fat dry milk for (2 hr, 20°C), washed in the same medium and incubated with the primary antibody (16 hr, 4°C) in the blocking buffer. After three washes, the membrane was incubated with the secondary antibody (2 hr, 20°C) and then washed. Protein bands were visualized using the Immobilon Forte Western HRP substrate (Millipore, WBLUF0500) in the Image Quant (LAS4000) imaging system. Densitometric quantification was performed on unsaturated images using ImageJ software. Antibodies used in this study are listed in Table 3

**Table 3:** Details of antibodies used.

| Antibody | Dilution | Catalog no. | Company |
| --- | --- | --- | --- |
| Beta-Tubulin | 1:5000 | Ab21058 | Abcam |
| Beclin1 (D40C5) | 1:1000 | 3495 | Cell Signalling Technology |
| LC3A/B (D3U4C) | 1:1000 | 12741 | Cell Signalling Technology |
| IP<sub>3</sub>R-II (A-5) | 1:1000 | sc-398434 | Santa Cruz Biotechnology |
| Anti-TRP2/DCT | 1:2000 | Ab74073 | Abcam |
| Anti-Melanoma GP100 | 1:2000 | Ab137078 | Abcam |
| Tyrosinase | 1:2000 | 35-6000 | Invitrogen |
| ULK1 (D8H5) | 1:1000 | 8054 | Cell Signalling Technology |
| Phospho-ULK1 (Ser757) | 1:1000 | 6888 | Cell Signalling Technology |
| AMPKα (D5A2) | 1:1000 | 5831 | Cell Signalling Technology |
| Phospho-AMPKα (Thr172) | 1:1000 | 2535 | Cell Signalling Technology |
| p70 S6 Kinase (49D7) | 1:1000 | 9202 | Cell Signalling Technology |
| Phospho-p70 S6 Kinase (Thr389) | 1:1000 | 97596 | Cell Signalling Technology |
| 4E-BP1 (53H11) | 1:1000 | 9644 | Cell Signalling Technology |
| Phospho-4E-BP1 (Thr37/46) (236B4) | 1:1000 | 2855 | Cell Signalling Technology |
| mTOR (7C10) | 1:1000 | 2983 | Cell Signalling Technology |
| Phospho-mTOR (Ser2448) (D9C2) | 1:1000 | 5536 | Cell Signalling Technology |
| TFEB | 1:1000 | 4240 | Cell Signalling Technology |
| Lamin | 1:1000 | MA5-35284 | Invitrogen |
| Beta-Actin | 1:10000 | A3854 | Sigma |

**Table 4:**
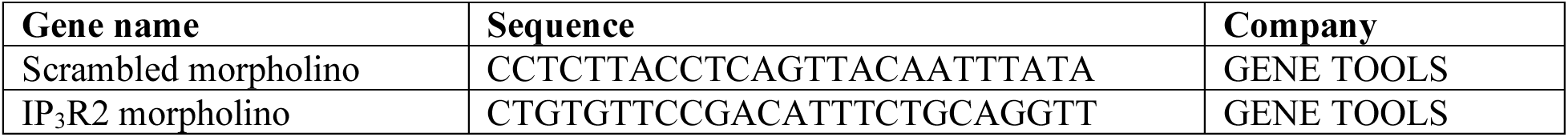
Details of morpholinos used:

### Melanin content assay

The melanin content of the samples was quantified using a previously described experimental protocol (Motiani *et al*, 2018; Tanwar *et al*, 2022a). Equal numbers of cells from each condition were lysed in 1 N sodium hydroxide solution through heating at 80°C for 3-4 hours. The samples were then centrifuged to remove any cellular debris. The absorbance of the resulting supernatants was measured at 405 nm using a Fluorescence Spectrophotometer. The melanin content of the samples was estimated by comparing their absorbance values to a standard curve (µg/ml) generated using synthetic melanin. In some samples, mean pixel intensity of cell pellet was calculated using image J (NIH).

### Cytosolic Ca^2+^ imaging

Intracellular Ca^2+^ levels were assessed through fluorescence imaging, as described in prior reports (Tanwar *et al*, 2022b). Cells were cultured on confocal dishes (SPL Life Sciences, 200350) and incubated with the ratiometric Ca^2+^ indicator fura-2AM (Thermo Fisher Scientific, F1221) for 30 minutes at 37°C. After washing the cells in Hepes-buffered saline solution and Ca^2+^-free HBSS buffer, specific cells were selected at 20X dry objective and imaged using a digital fluorescence microscopy system (Nikon Eclipse Ti2 microscope equipped with a CoolLED pE-340 Fura light source and a high-speed PCO camera). Fura-2AM was alternately excited at 340 and 380 nm, and the corresponding emission signals were captured at 510 nm. The resulting Ca^2+^ traces represent the average response from multiple cells within a single imaging dish, with the number of cells (n) indicated for each trace.

### Mitochondrial Ca^2+^ imaging

Intracellular mitochondrial Ca^2+^ levels were measured using fluorescence-based Ca^2+^ imaging as described previously (Tanwar *et al*, 2024). B16 cells were seeded on confocal dishes. After the required treatment and transfections, 1.5µg of pCMV CEPIA2mt plasmid (a gift from Masamitsu lino, Addgene, 58218) per dish was transfected using TurboFect at a 1:2 ratio (w/v). The following day, the cells were washed three times with Ca^2+^-free HEPES-buffered saline solution and incubated in HEPES-buffered saline solution. Several cells were selected using a 60X oil objective, and imaging was performed using a digital fluorescence microscopy system (Nikon Eclipse Ti2 microscope equipped with a CoolLED pE-340 Fura light source and a high-speed PCO camera). The CEPIA2mt-transfected cells were excited at 488 nm, and the emission signal was captured at 500-550 nm. 50 µM histamine was used as a stimulus to release Ca^2+^ from the IP_3_ receptor channels, and the resulting changes in mitochondrial Ca^2+^ were measured using the CEPIA2mt probe.

B16-F10 cells were cultured in cover-glass bottom dishes to study the functional role of IP_3_R2. Rescue experiments were conducted by overexpressing either wild-type mouse IP_3_R2 or a mutant variant (a gift from Dr. David Yule) in cells with stable IP_3_R2 knockdown. Cultured B16 cells (IP_3_R^2+/+^, IP_3_R2 ^-/-^) were seeded in cover-glass bottom dishes and transfected with 1.5µg of the CEPIA2mt Ca^2+^ indicator construct, along with 1µg of either the mouse IP_3_R2 or the mouse IP_3_R2 mutant plasmid, using a 1:2 ratio (w/v) of TurboFect transfection reagent at 50-60% confluency. After 48 hours, several transfected cells were selected, and intracellular Ca^2+^ dynamics were examined using fluorescence-based Ca^2+^ imaging as described previously.

### TRPML1-G-GECO1.2-ERES Ca^2+^ imaging

Lysosomal Ca^2+^ measurements were conducted as previously described (Davis *et al*, 2020). Following the necessary transfections, B16 cells were transfected with 1.5µg of the TRPML1-G-GECO1.2-ERES (Addgene, 207144) construct at 50-60% confluency in cover-glass bottom dishes. The cells were washed three times with Ca2+-free HBSS buffer the next day. Specific cells were selected, and imaging was performed using a digital fluorescence microscopy system with a 60X oil objective (Nikon Eclipse Ti2 microscope equipped with a CoolLED pE-340 Fura light source and a high-speed PCO camera). The TRPML1-G-GECO1.2-ERES transfected cells were excited at 488 nm, and the corresponding emission signals were captured at 500-550 nm. 20 µM MLSA1(Sigma Aldrich., SML0627) or 300 µM GPN (Gly-Phe β-naphthylamide, sc-252858) was utilized as a stimulus to induce Ca^2+^ release from the TRPMLs channels. The resulting Ca^2+^ traces represent the average response from multiple cells within a single imaging dish, with the number of cells (n) indicated for each trace.

B16-F10 cells were cultured in cover-glass bottom dishes to study the functional role of Mucolipin1 or TRPML1. The cells were co-transfected with 1.5 µg of TRPML1-G-GECO1.2-ERES and 1 µg of Mucolipin1-pHcRed C1 (Addgene, 62959) or the TRPML1 pore-dead mutant, Mucolipin1 D471-472K-pHcRed C1 (Addgene, 62961), at 50-60% confluency using TurboFECT transfection reagent at a 1:2 ratio (w/v). Forty-eight hours post-transfection, the cells were washed three times with Ca^2+^-free 1X HBSS buffer. Green fluorescent cells were selected, and Ca^2+^ imaging was performed using a digital fluorescence microscopy system with a 60X oil objective. 20 µM MLSA1 as the TRPML channel agonist was used as a stimulus to induce Ca^2+^ release.

### Oregon green 488 BAPTA-1 dextran (OBDx)

Lysosomes of the cells were identified through the endocytosis of a Ca^2+^-sensitive probe, as previously described (Kim *et al*, 2021). Specifically, the cells were incubated with 100 µg/mL of Oregon-BAPTA-dextran (Thermo Fisher Scientific, O6798) in the culture medium for 12 hours, followed by an additional 12-hour pulse-chase period with phenol red-free DMEM to allow for lysosomal staining. The cells were then washed thrice with HBSS buffer. The intensity of the Ca^2+^-dependent fluorescence was measured using Confocal microscopy equipped with a 63X/1.40 oil immersion objective (Laser Scanning Confocal Microscope: LSM 880, Carl Zeiss). Cells were excited with a 488 nm laser, and the resulting fluorescence emission was captured within the 497-572 nm range. The Ca^2+^ fluorescence intensity was quantified using Fiji (NIH) software.

### pMRX-IP-GFP-LC3-RFP plasmid transfection

Autophagy flux was analyzed using fluorescence microscopy by monitoring the degradation of the GFP-LC3 reporter, with RFP serving as an internal control, as previously described (Kaizuka *et al*, 2016). Cultured B16 cells seeded in cover-glass bottom dishes were transfected with the pMRX-IP-GFP-LC3-RFP plasmid (Addgene, 84573) using Turbofect Transfection reagent at a 1:2 ratio (w/v) for 48 hours. Live cell images were acquired using confocal microscopy equipped with a 63X/1.40 oil immersion objective (Laser Scanning Confocal Microscope: LSM 880, Carl Zeiss). Cells were excited with 543 nm and 488 nm lasers in an alternating manner, and the resulting fluorescence emissions were captured at 551-632 nm and 497-572 nm ranges, respectively. The confocal images were then deconvoluted using ZEN 2.3 SP1 FP1 (black) software (version 14.0) and analyzed using Imaris software.

### Cloning and transfection of mCherry-Tyrosinase-EGFP plasmid

The mCherry-Tyrosinase-EGFP melanophagy construct consists of the Tyrosinase sequence targeting specifically to Melanosome. To clone mCherry, a red fluorescent protein to pEGFP Tyrosinase (Addgene, 32781), mCherry was amplified from vector mCherry N1(Clontech, 632523) having NHE1 and XHO1 sites present at 633-1386bp. Amplified Vector mcherry N1 cloned upstream of Tyrosinase-eGFP of NHE1-XHO1 site at 591-613bp of eGFP Tyrosinase plasmid construct to generate RFP-Tyrosinase-GFP.

Melanophagy flux was analyzed by examining the differential stability of the green and red fluorescent proteins within the acidic lysosomal environment. B16 cells were transfected with the cloned RFP-Tyrosinase-GFP plasmid construct at 50-60% confluency using a TurboFect transfection reagent at a 1:2 ratio (w/v). Twenty-four hours post-plasmid transfection, live-cell images were acquired using confocal microscopy equipped with a 63X/1.40 oil immersion objective (Laser Scanning Confocal Microscope: LSM 880, Carl Zeiss). Cells were alternatingly excited with 543 nm and 488 nm lasers, and the resulting fluorescence emissions were captured at 551-632 nm and 497-572 nm ranges, respectively. The confocal images were then deconvoluted using ZEN 2.3 SP1 FP1 (black) software (version 14.0) and analyzed using Imaris software.

### Cloning and transfection of Tyrosinase-mKeimaN1 plasmid

Keima is a fluorescent protein derived from coral that exhibits pH-dependent emission spectra, emitting distinct colors in acidic and neutral environments. The cumulative fluorescence signal can be used to quantify autophagy at a single time point. The Tyrosinase-mKeimaN1 melanophagy construct consists of the Tyrosinase sequence specifically targeting Melanosomes. To generate this construct, the Tyrosinase sequence was amplified from the eGFP-TYR plasmid (Addgene, 32781), which harbors ECOR1 and KPN1 restriction sites at 629-2231 base pairs. The amplified Tyrosinase sequence was then cloned upstream of the mKeima-Red-N1 vector (Addgene,54597), between the ECOR1 and KPN1 sites at 665-685 base pairs, to create the Tyrosinase-mKeima melanophagy reporter construct.

B16 cells were cultured and transfected with 2.5 µg of the Tyrosinase-mKeima N1 construct using TurboFECT transfection reagent at a 1:2 ratio (w/v) when the cells were 50-60% confluent. After 48 hours of plasmid transfection, live-cell imaging was performed using a confocal microscope equipped with a 63X/1.40 oil immersion objective (Laser Scanning Confocal Microscope: LSM 880, Carl Zeiss). Cells were excited with 543 nm and 488 nm lasers in an alternating manner, and the resulting fluorescence emissions were captured in the 556-678 nm and 493-620 nm ranges, respectively. The confocal images were then deconvoluted using ZEN 2.3 SP1 FP1(black) software (version 14.0) and analyzed using Imaris software.

### Measuring the distance between ER and Mitochondria

A split-GFP-based contact site sensor (SPLICS) has been developed to quantify the contact sites between the endoplasmic reticulum and mitochondria across a range of distances. B16 cells were transfected with 1µg of the SPLICS Mt-ER Short P2A plasmid (Addgene, 164108) using the TurboFect transfection reagent at a 1:2 ratio (w/v). Live-cell imaging was performed 24 hours after the plasmid transfection, utilizing a confocal microscope equipped with a 63X/1.40 oil immersion objective (Laser Scanning Confocal Microscope: LSM 880, Carl Zeiss). The cells were excited with 488 nm lasers, and the resulting fluorescence emissions were captured within the 497-572 nm range. The confocal images were then deconvoluted using ZEN 2.3 SP1 FP1 (black) software (version 14.0) and analyzed using Fiji software.

### Cycloheximide chase assay

B16 cells at 50-60% confluency were transfected with siRNA using TurboFect transfection reagent at 1:3 ratio. 1µM Melanocyte Stimulating Hormone was added 24 hours post-transfection. 72 hours after transfection, the cells were incubated with 20µg/ml cycloheximide (Abcam, 120093) for 0, 4, and 8 hours. Subsequently, the cells were harvested, lysed, and prepared for western blotting analysis.

### Measuring Lysosomal pH

To assess changes in lysosomal pH, B16 cells grown on cover-glass bottom dishes were stained with the Lysosensor Yellow-Blue DND-160 dye at a 1:500 ratio in growth medium for 5 minutes in a humidified CO2 incubator at 37°C. The cells were then washed thrice with 1X HBSS buffer, and live-cell imaging was performed using a confocal microscope equipped with a 63X/1.40 oil immersion objective (Laser Scanning Confocal Microscope: LSM 880, Carl Zeiss). The Lysosensor Yellow-Blue imaging was conducted by exciting the samples at 405 nm and capturing the emission signals in the 490-556 nm (yellow in more acidic organelles) and 417-481 nm (blue in less acidic organelles) ranges (Prajapat *et al*, 2024). The confocal images were deconvoluted using ZEN 2.3 SP1 FP1 (black) software (version 14.0) and analyzed using Imaris software.

pHluorin2 is an advanced, ratiometric, pH-sensitive green fluorescent protein. This GFP variant exhibits a dual-modal excitation spectrum, whereby acidification leads to a decrease in 405 nm excitation coupled with a corresponding increase in 488 nm excitation. Cells cultured on cover-glass bottom dishes were transfected with 0.5 µg of LAMP1-RpHLuorin2 plasmid (Addgene, 171720) using Turbofect Transfection reagent at a 1:2 ratio (w/v). After 24 hours post-transfection, live-cell imaging was performed using a confocal microscope equipped with a 63X/1.40 oil immersion objective (Laser Scanning Confocal Microscope: LSM 880, Carl Zeiss). The cells were alternately excited with 405 nm and 488 nm lasers, and the resulting fluorescence emissions were captured within the 490-560 nm range. The confocal images were then deconvoluted using ZEN 2.3 SP1 FP1 (black) software (version 14.0) and analyzed using Imaris software.

### TFEB nuclear translocation

B16 melanoma cells cultured in cover-glass bottom dishes at 50-60% confluency were transfected with small interfering RNA (siRNA) using the TurboFect transfection reagent at a 1:3 ratio. After 24 hours, 1µM Melanocyte Stimulating Hormone was added to the cells. Forty-eight hours post-siRNA transfection, the B16 cells were transfected with 1µg of the TFEB-GFP plasmid construct using the TurboFECT reagent at a 1:2 ratio (w/v). Before live-cell confocal microscopy, the cells were stimulated with 20 µM MLSA1 for 1 hour. The confocal imaging was performed using a confocal microscope equipped with 63X/1.40 oil immersion objective (Laser Scanning Confocal Microscope: LSM 880, Carl Zeiss). The cells were incubated with Hoechst, a nuclear stain, at a 1:2000 dilution for 5 minutes to label the nuclei. The cells were then excited with 405 nm and 488 nm lasers alternatingly, and the resulting fluorescence emissions were captured within the 420-486 nm and 510-587 nm wavelength ranges, respectively. The acquired confocal images were then deconvoluted using the ZEN 2.3 SP1 FP1 (black) software (version 14.0) and analyzed using the Fiji software (NIH).

### Subcellular fractionation

The cells were collected and resuspended in a cytosolic buffer (1X PBS, 0.1% NP-40, 1X Protease Inhibitor Cocktail). This mixture was incubated for 10 minutes at room temperature, representing the whole-cell lysate. The lysates were then centrifuged at 10,000 rpm for 1 minute. The supernatant obtained from this step constitutes the cytoplasmic extract. The pellet was resuspended in 4X loading dye (100 mM Tris-HCl, 4% SDS, 20% glycerol, 0.2% bromophenol) and heated at 95°C for 20-30 minutes.

### Proximity ligation assay

B16 cells at 50-60% confluency were transfected with siRNA using a TurboFect transfection reagent at 1:3. 1µM Melanocyte Stimulating Hormone was added 24 hours post-transfection. At 72 hours after transfection, cells grown on a coverslip were fixed using 4% Paraformaldehyde (PFA) at room temperature for 15 mins. The cells were then washed with 1x PBS and permeabilized using 0.3% TritonX-100 for 15 mins at room temperature. To investigate ER-lysosome interactions, primary antibodies against VAP-A (ER) (Santa Cruz Biotechnology, sc-293278) and LAMP1 (lysosomes) (Cell signaling Technology, 9091) were used. Incubations with Duolink PLA probe anti-rabbit PLUS (Sigma, DUO92002) and anti-mouse MINUS (Sigma, DUO92004), ligase and polymerase (Sigma, DUO92013), and the washes between each step were precisely as recommended by the manufacturer’s protocol. Cells were then mounted in SlowFade™ Gold Antifade Mountant with DAPI (Invitrogen, S36938) to label the nucleus. PLA products were visualized using the Zeiss microscope with 63X, and spots were quantified using Fiji software.

### ADP/ATP measurement assay

B16 cells at 50-60% confluency were transfected with siRNA using a TurboFect transfection reagent at 1:3. 1µM Melanocyte Stimulating Hormone was added 24 hours post-transfection. 72 hours post-transfection, cells were trypsinized and followed the manufacturer’s protocol (Sigma, MAK135). Data was acquired using a Luminometer and ratio was calculated according to the manufacturer’s protocol.

### Transmission Electron Microscopy (TEM)

Cells were collected and resuspended in Phosphate buffer (PB). Remove PB buffer by centrifugation, and cells were resuspended in TEM fixative for 2hrs at 4℃. Remove the TEM fixative by centrifugation and resuspend it in the PB buffer. Samples were processed, and ultrathin sections were sliced at around 60-90nm. The double staining method uses uranyl acetate and alkaline lead citrate to obtain a good contrast of sections (Reynolds, 1963). TEM images captured in JEM1400 Flash equipped with tungsten filament as an electron source and a highly sensitive sCMOS camera. Images captured at a magnification of 30,000X and a high voltage of 80kV.

### Zebrafish husbandry

Zebrafish used in this study were housed in the Laboratory of Calciomics and Systemic Pathophysiology at the Regional Centre for Biotechnology, India, with proper standard ethical protocols approved by the Institutional Animal Ethics Committee of Regional Centre for Biotechnology, India, with care to minimize animal suffering.

### Morpholino design and microinjections

ATG blocking antisense morpholino oligos were designed against zebrafish IP_3_R2 gene according to manufactures protocol (GENE TOOLS).

### Overexpression and complementation/rescue assay of IP_3_R2 in zebrafish

Plasmids were linearized using not1 and then used as a template for generating an In-vitro transcript using the mMESSAGE mMACHINE T7 Transcription Kit according to the manufacturer’s protocol.

### Statistical analysis

All experiments were performed at least 3 times. All statistical analysis was performed using Graph pad prism8.0 software. Data are represented as mean ± SEM. An unpaired student t-test or one sample t-test was performed to check statistical significance between 2 groups. A one-way ANOVA test was performed to compare the mean of more than 2 groups. Two-way ANOVA was performed to compare the mean between two groups at different time points. p-value < 0.05 was considered as significant and presented as ‘’*’’, p-value < 0.01 is presented as “**”, p-value < 0.001 is presented as “***” and p-value < 0.0001 is presented as “****”.

## Authors Contribution

Suman Saurav: Methodology, Investigation, Visualization, Formal analysis, Writing-Original draft preparation. Rajender K Motiani: Conceptualization, Supervision, Writing-Original draft preparation, Reviewing and Editing, Project administration, Funding acquisition.

## Competing interests

Authors declare that they have no competing interests.

## Acknowledgements

This work was supported by the DBT/Wellcome Trust India Alliance Fellowship (IA/I/19/2/504651). RKM also acknowledges funding support from RCB Institutional Core funding. We thank Dr. David I. Yule (University of Rochester, USA) for sharing IP_3_R2 and IP_3_R2-M plasmids. We also thank Dr. Manjula Kalia (Regional Centre for Biotechnology, India) for sharing reagents and antibodies. The authors thank members of the Motiani laboratory for the critical discussions. SS acknowledges his Junior and Senior Research Fellowship from CSIR, India.

---

**Supplementary Figure 1:**
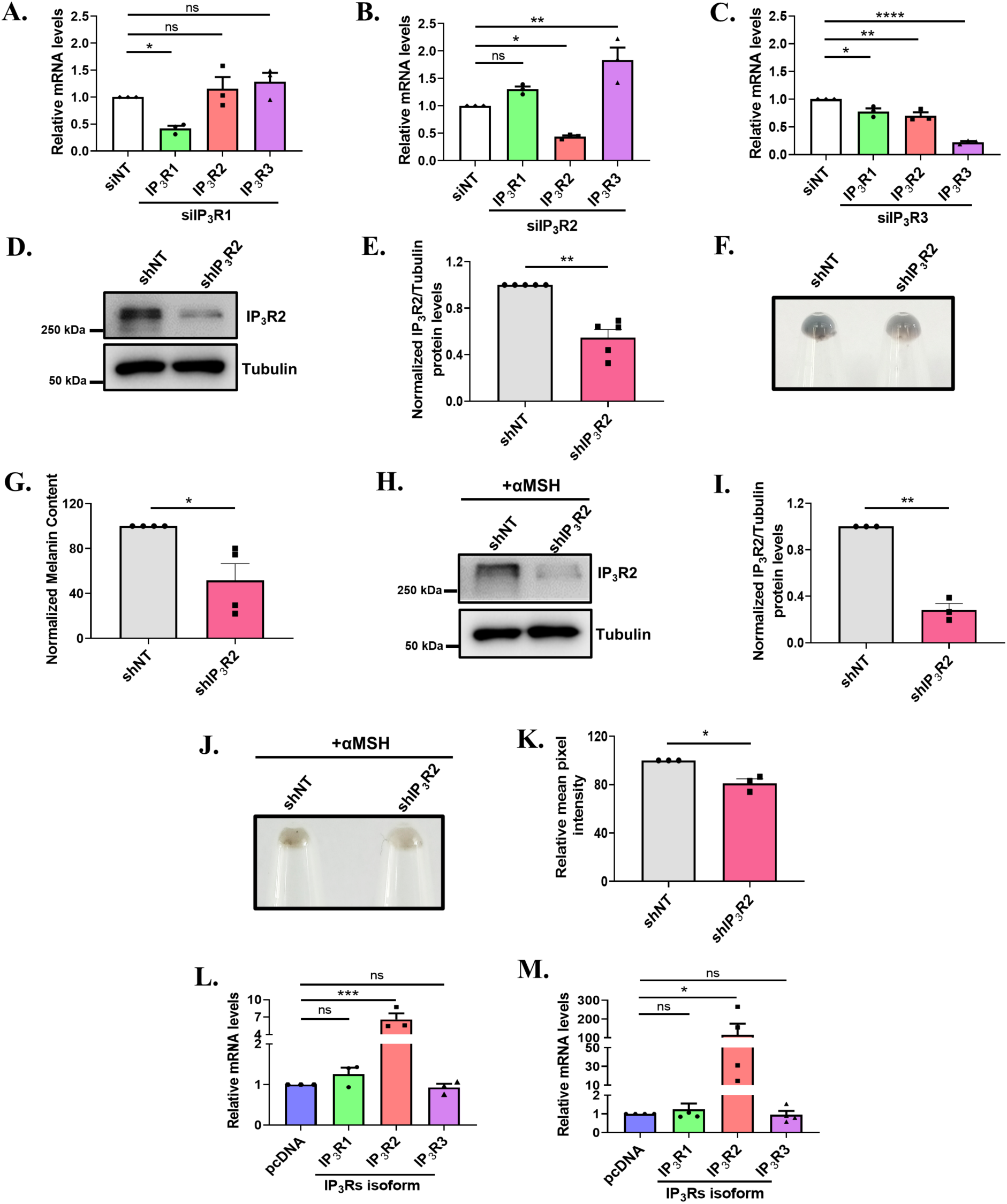
IP_3_R2 positively regulates pigmentation. (A) qRT PCR analysis showing expression of IP_3_Rs isoform after 72hrs of siRNA mediated IP_3_R1 silencing in LD model system (N=3). (B) qRT PCR analysis showing expression of IP_3_Rs isoform after 72hrs of siRNA mediated IP_3_R2 silencing in LD model system (N=3). (C) qRT PCR analysis showing expression of IP_3_Rs isoform after 72hrs of siRNA mediated IP_3_R3 silencing in LD model system (N=3). (D) Representative image of immunoblot showing expression of IP_3_R2 in B16 cells after stable knockdown of IP_3_R2 in LD model system. (E) Bar graph showing the densitometry of IP_3_R2 band normalized to β-Tubulin (N=3). (F) Representative image of pellet pictures shows B16 cells having stable knockdown of IP_3_R2 in LD model system. (G) Bar graph showing mean pixel intensity of B16 cells having stable knockdown of IP_3_R2 in LD model system (N=3). (H) Representative image of immunoblot showing expression of IP_3_R2 in B16 cells after stable knockdown of IP_3_R2 with 48hrs exposure to 1µM of αMSH. (I) Bar graph showing the densitometry of IP_3_R2 band normalized to β-Tubulin (N=3). (J) Representative image of pellet pictures shows B16 cells having stable knockdown of IP_3_R2 along with adding 1µM αMSH. (K) Bar graph showing mean pixel intensity of B16 cells having stable knockdown of IP_3_R2 along with adding 1µM αMSH (N=3). (L) qRT PCR analysis showing expression of IP_3_Rs isoform after 72hrs of overexpression with functional IP_3_R2 in LD model system (N=3). (M) qRT PCR analysis showing expression of IP_3_Rs isoform after 72hrs of overexpression with functional IP_3_R2-M in LD model system (N=4). Data presented are mean ± SEM. For statistical analysis, one sample t-test was performed for panels E, G, I, K and Dunnett’s multiple comparisons test was performed for panels A, B, C, L, M using GraphPad Prism software. Here, ‘ns’ means non-significant; * p <0.05, ** p < 0.01, *** p < 0.001and **** p < 0.0001.

**Supplementary Figure 2:**
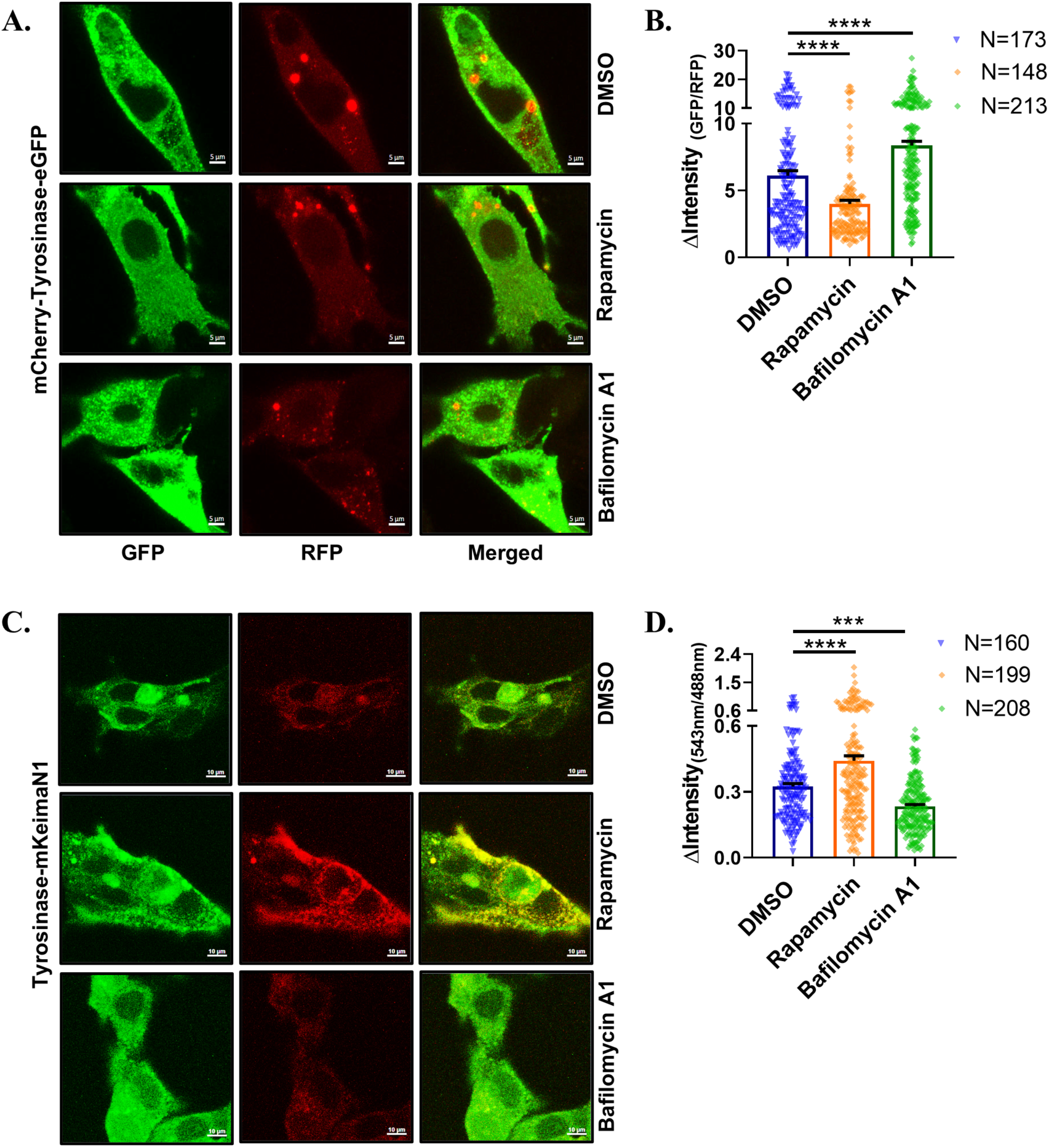
Functional validation of melanophagy probes. (A) Representative images of confocal imaging in B16 cells stimulated with 4µM rapamycin and 100nM BafilomycinA1 for 5hrs after transfection with mCherry-Tyrosinase-EGFP probe, scale bar, 5µm. (B) Bar graph shows the quantification of the autolysosome intensity in the GFP:RFP ratio, where ‘N’ denotes the number of cells. (C) Representative images of confocal imaging in B16 cells stimulated with 4µM rapamycin and 100nM BafilomycinA1 for 5hrs after transfection with Tyrosinase-mKeimaN1 probe, scale bar, 10µm (D) Bar graph shows the quantification of the autolysosome intensity in the 543nm:488nm ratio, which is alternatively excited at 543nm and 488nm, where ‘N’ denotes the number of cells. Data presented are mean ± SEM. For statistical analysis, Dunnett’s multiple comparisons test was performed for panels B, D using GraphPad Prism software. Here, *** p < 0.001 and **** p < 0.0001.

**Supplementary Figure 3:**
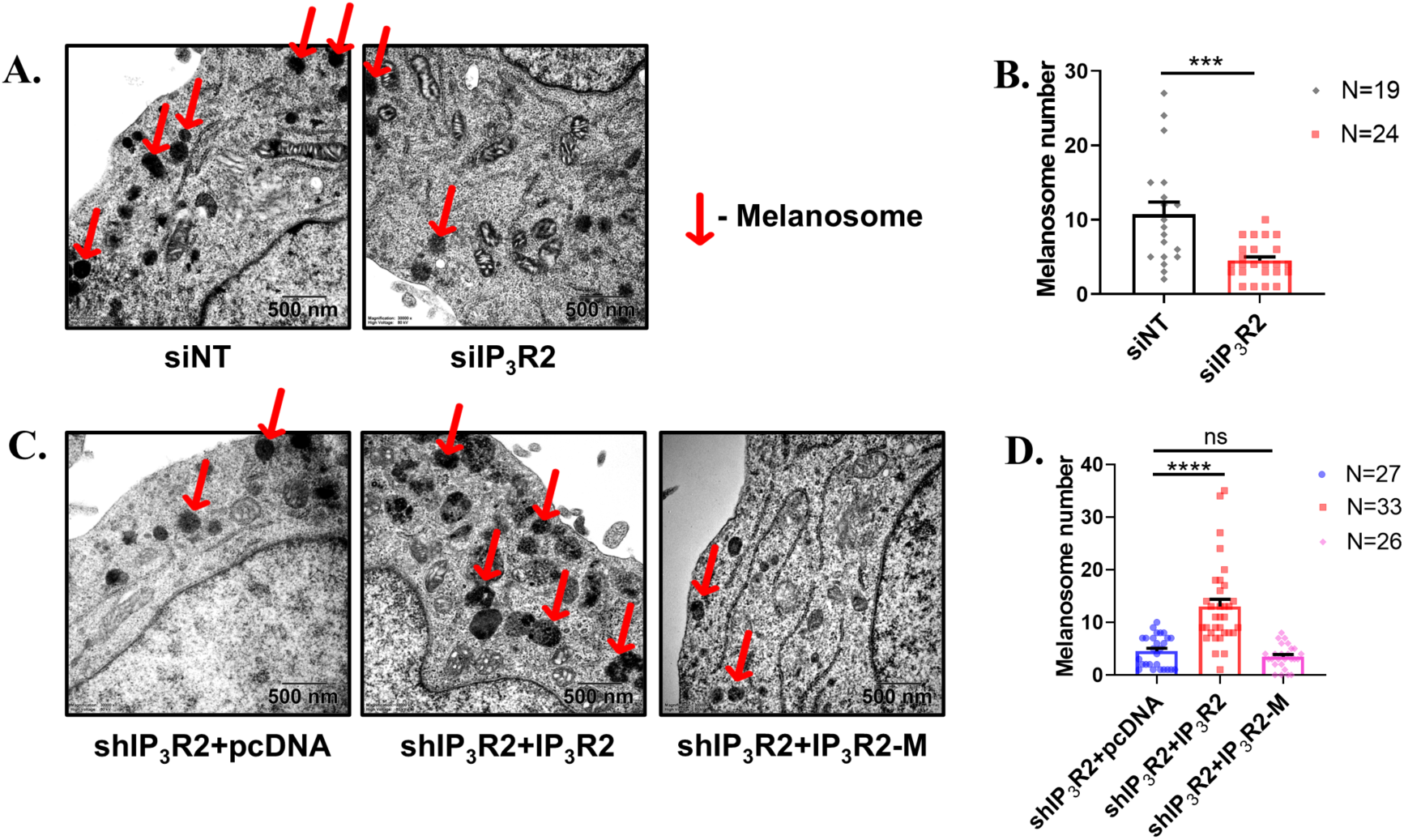
IP_3_R2 silencing leads to melanosome degradation. (A) Representative TEM image showing B16 cells transfected with siNT or siIP_3_R2, scale bar, 500nm. (B) Bar graph shows the quantification of the melanosome number, where ‘N’ denotes the melanosome number per image. (C) Representative TEM image showing B16 cells having stable IP_3_R2 knockdown demonstrating rescue with IP_3_R2 and IP_3_R2-M, scale bar, 500nm. (D) Bar graph shows the quantification of the melanosome number, where ‘N’ denotes the melanosome number per image. Data presented are mean ± SEM. For statistical analysis, unpaired t-test was performed for panel B and Dunnett’s multiple comparisons test was performed for panel D, using GraphPad Prism software. Here, ‘ns’ means non-significant; ** p < 0.01, *** p < 0.001and **** p < 0.0001.

**Supplementary Figure 4:**
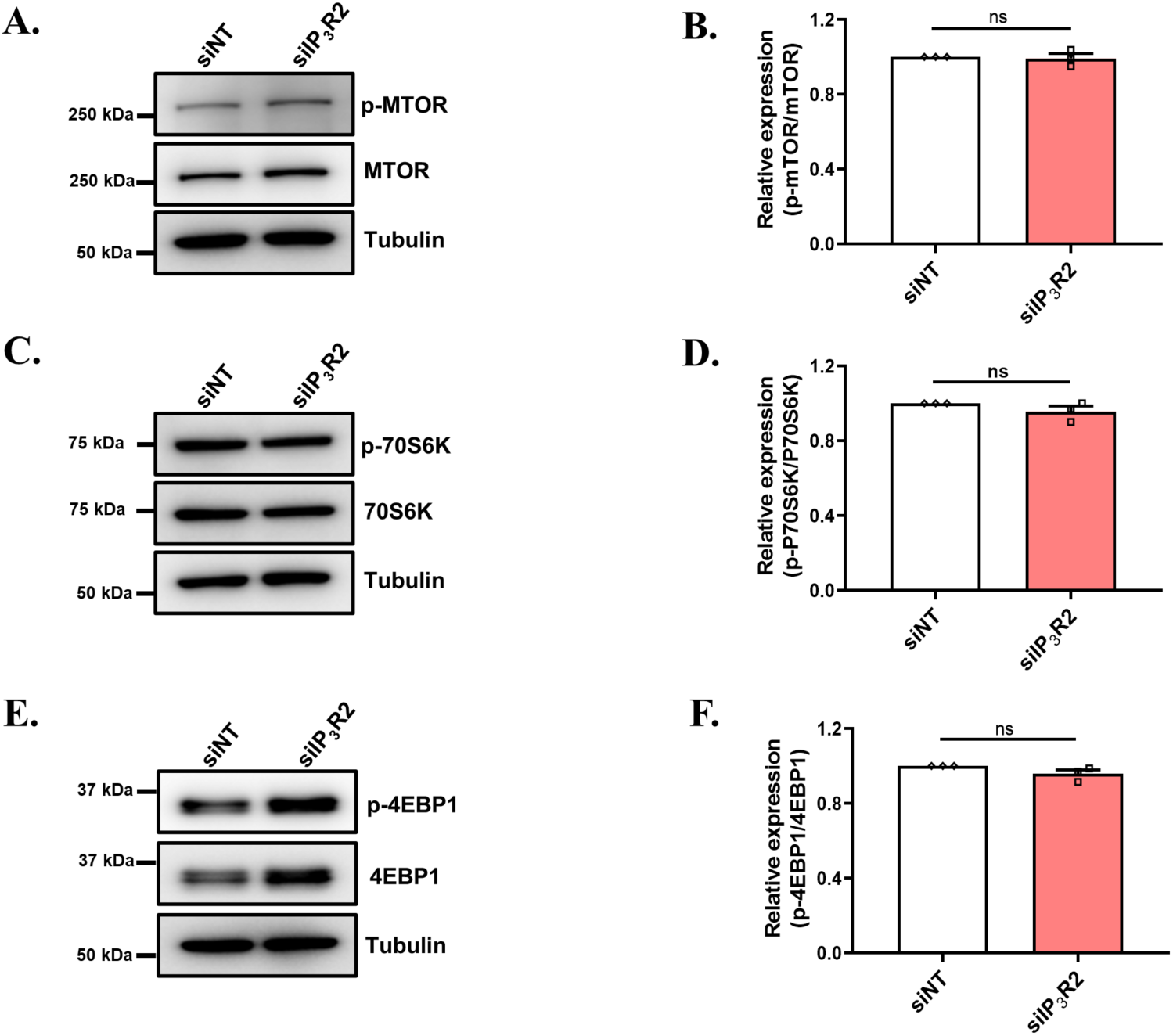
IP_3_R2 knockdown-induced melanophagy is mTOR independent. (A) Representative image of immunoblot showing expression of p-mTOR and total mTOR in B16 cells after siRNA silencing of IP_3_R2. (B) Bar graph showing the densitometric analysis of the relative changes in the levels of p-mTOR to total mTOR (N=3). (C) Representative image of immunoblot showing expression of p-70S6K and total 70S6K in B16 cells after siRNA silencing of IP_3_R2. (D) Bar graph showing the densitometric analysis of the relative changes in the levels p-70S6K to total 70S6K (N=3). (E) Representative image of immunoblot showing expression of p-4EBP1 and total 4EBP1 in B16 cells after siRNA silencing of IP_3_R2. (F) Bar graph showing the densitometric analysis of the relative changes in the p-4EBP1 to total 4EBP1 (N=3). Data presented are mean ± SEM. For statistical analysis, one sample t-test was performed for panels B, D, F, using GraphPad Prism software. Here, ‘ns’ means non-significant; ** p < 0.01.

**Supplementary Figure 5:**
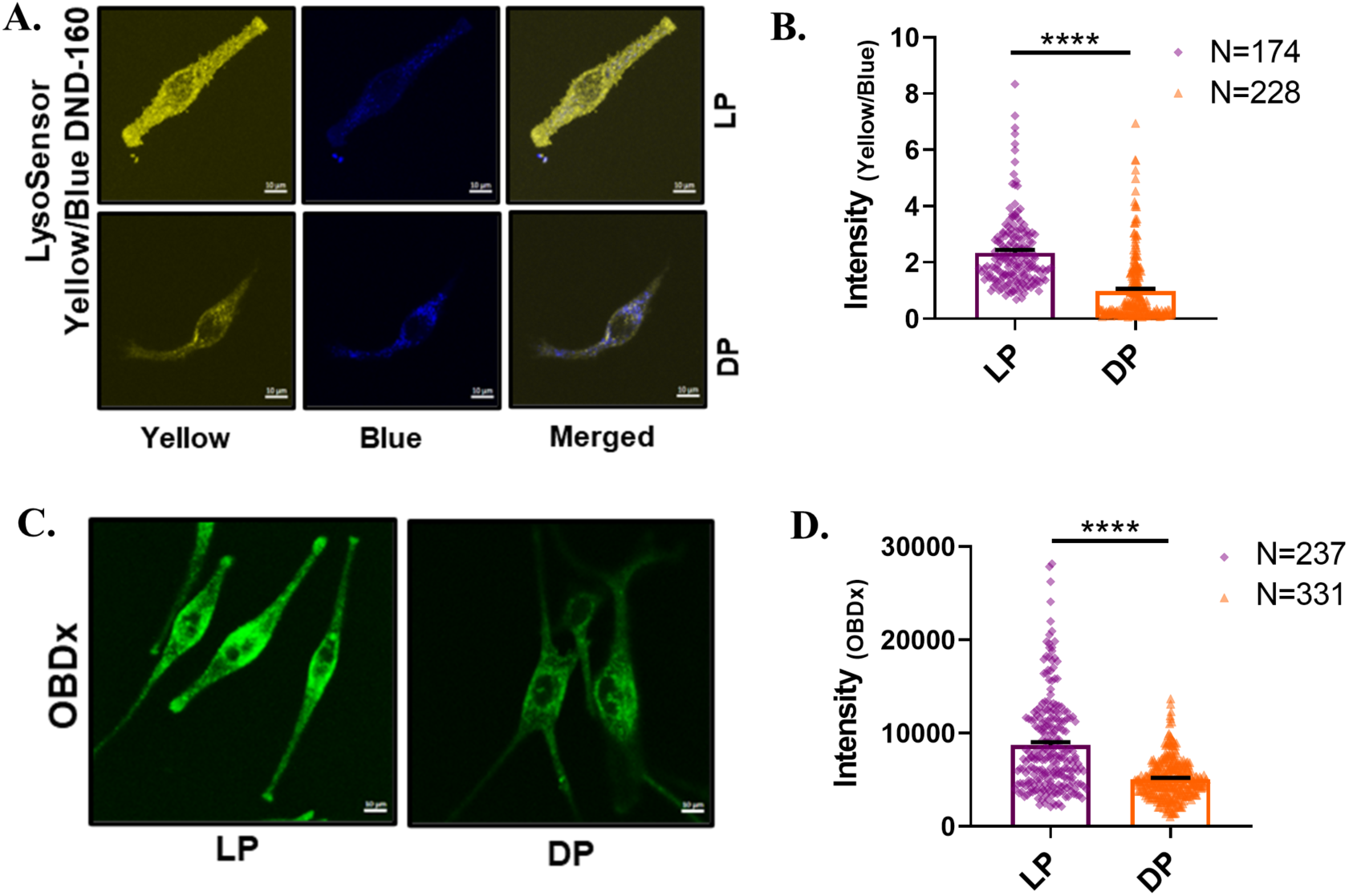
Lightly pigmented melanocytes have lower lysosomal pH and higher lysosomal calcium levels in comparison to darkly pigmented primary melanocytes. (A) Representative images of confocal imaging in B16 cells loaded with LysoSensor Yellow/Blue DND-160 dye for 5 mins in lightly-pigmented and darkly pigmented primary melanocytes, scale bar, 10µm. (B) Bar graph showing the quantification of the intraluminal lysosomal pH in Yellow/Blue ratio, where ‘N’ denotes the number of cells imaged. (C) Representative images of confocal imaging in B16 cells loaded with OG-BAPTA-dextran (OBDx) in lightly-pigmented and darkly pigmented primary melanocytes, scale bar, 10µm. (D) Bar graph showing the quantification of the GFP intensity, where ‘N’ denotes the number of cells imaged. Data presented are mean ± SEM. For statistical analysis, an unpaired t-test was performed for panels B and D using GraphPad Prism software. Here, **** p < 0.0001.

**Supplementary Figure 6:**
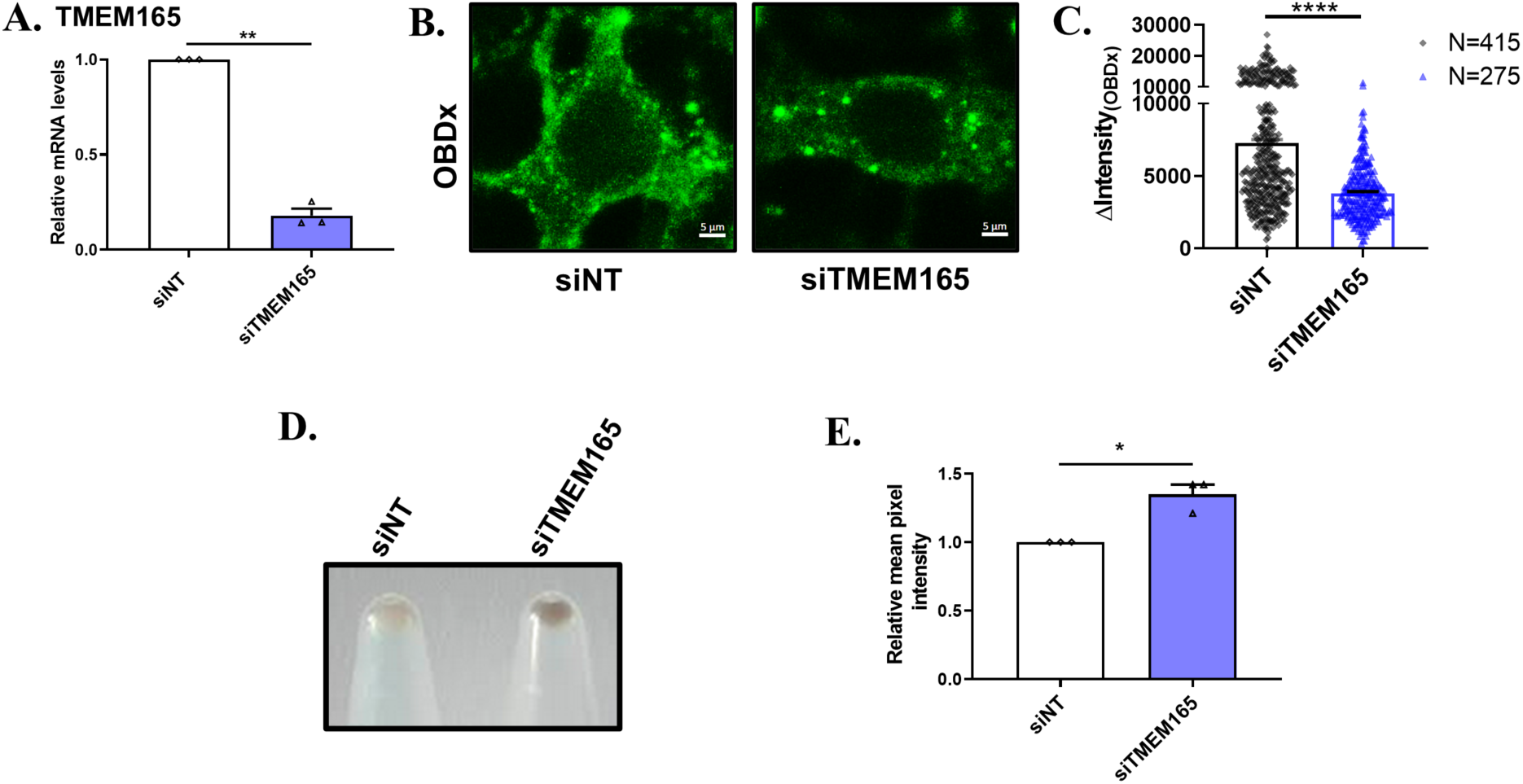
TMEM165 knockdown decreases lysosomal calcium, thereby enhancing pigmentation. (A) qRT PCR analysis showing decrease expression of TMEM165 after 72hrs of siRNA mediated TMEM165 silencing. (B) Representative images of confocal imaging in B16 cells loaded with OG-BAPTA-dextran (OBDx) after transfected with siNT or siTMEM165, scale bar, 5µm. (C) (H) Bar graph showing the quantification of the GFP intensity, where ‘N’ denotes the number of cells imaged. (D) Representative image of pellet pictures shows B16 cells transfected with siNT or siTMEM165. (E) Bar graph showing mean pixel intensity of B16 cells transfected with siNT or siTMEM165 (N=3). Data presented are mean ± SEM. For statistical analysis, an unpaired t-test was performed for panel C and one sample t-test was performed for panels A, and E using GraphPad Prism software. Here, * p <0.05, ** p < 0.01, and **** p < 0.0001.

**Supplementary Figure 7:**
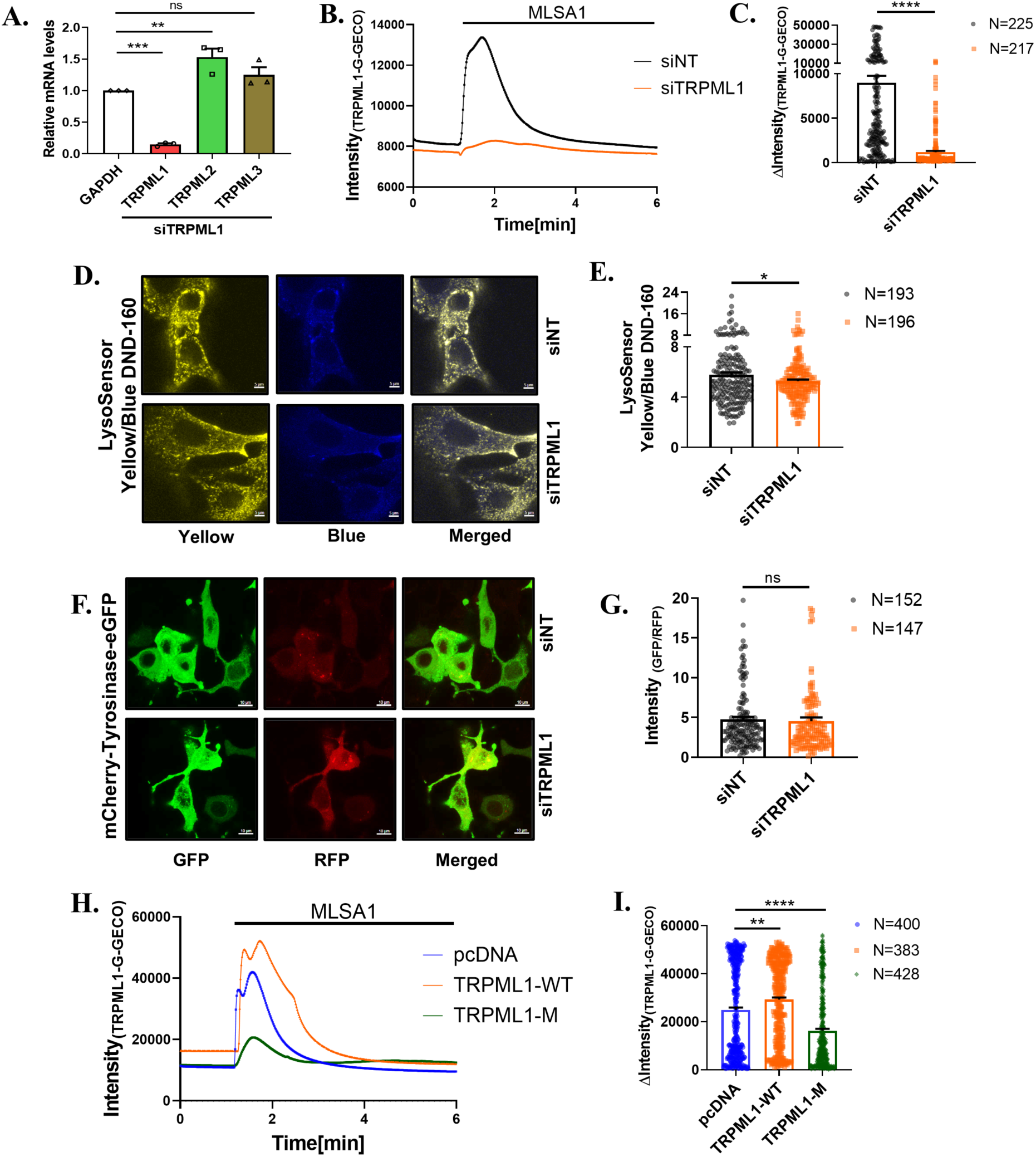
IP_3_R2 knockdown enhances TRPML1 activity. (A) qRT PCR analysis showing expression of TRPMLs isoform after 48hrs of siRNA mediated TRPML1 silencing (N=3). (B) Representative Ca^2+^ imaging trace using the TRPML1-G-GECO probe in B16 cells stimulated with the MLSA1 after 72hrs of siRNA transfection. (C) Bar graph showing the quantification of Ca^2+^ imaging traces stimulated with 20µM MLSA1, where ‘N’ denotes the total number of ROI in that trace. (D) Representative images of confocal imaging in B16 cells loaded with LysoSensor Yellow/Blue DND-160 dye for 5 mins after transfected with siNT or siTRPML1, scale bar, 5µm. (E) Bar graph showing the quantification of the intraluminal lysosomal pH in Yellow/Blue ratio, where ‘N’ denotes the number of cells imaged. (F) Representative images of confocal imaging in B16 cells after transfected with siNT or siTRPML1 along with mCherry-Tyrosinase-EGFP probe, scale bar, 10µm (G) Bar graph shows the quantification of the autolysosome intensity in the GFP:RFP ratio, where ‘N’ denotes the number of cells (H) Representative Ca^2+^ imaging trace using the TRPML1-G-GECO probe in B16 cells stimulated with the MLSA1 after overexpressing with TRPML1-WT and TRPML1-M. (I) Bar graph showing the quantification of Ca^2+^ imaging traces stimulated with 20µM MLSA1, where ‘N’ denotes the total number of ROI in that trace. Data presented are mean ± SEM. For statistical analysis, Dunnett’s multiple comparisons test was performed for panels A, I, and an unpaired t-test was performed for panels C, E, G using GraphPad Prism software. Here, ‘ns’ means non-significant; * p <0.05, ** p < 0.01, *** p < 0.001and **** p < 0.0001.

## Notes

### Competing Interest Statement

The authors have declared no competing interest.

## References

Ahuja K, Raju S, Dahiya S & Motiani RK (2025) ROS and calcium signaling are critical determinant of skin pigmentation. Cell Calcium 125: 102987

Antonia RJ & Baldwin AS (2018) IKK promotes cytokine-induced and cancer-associated AMPK activity and attenuates phenformin-induced cell death in LKB1-deficient cells. Sci Signal 11: 1–11

Arora S, Tanwar J, Sharma N, Saurav S & Motiani RK (2021) Orai3 Regulates Pancreatic Cancer Metastasis by Encoding a Functional Store Operated Calcium Entry Channel. Cancers (Basel) 13

Atakpa P, Thillaiappan NB, Mataragka S, Prole DL & Taylor CW (2018) IP3 Receptors Preferentially Associate with ER-Lysosome Contact Sites and Selectively Deliver Ca2+ to Lysosomes. Cell Rep 25: 3180–3193.e7

Bartok A, Weaver D, Golenár T, Nichtova Z, Katona M, Bánsághi S, Alzayady KJ, Thomas VK, Ando H, Mikoshiba K, et al (2019) IP3 receptor isoforms differently regulate ER-mitochondrial contacts and local calcium transfer. Nat Commun 10: 1–14

Bellono NW, Kamme LG, Zimmerman AL & Oancea E (2013) UV light phototransduction activates transient receptor potential A1 ion channels in human melanocytes. Proc Natl Acad Sci U S A 110: 2383–2388

Bellono NW & Oancea E V. (2014) Ion transport in pigmentation. Arch Biochem Biophys 563: 35–41

Buffey JA, Edgecombe M & Neil S MAC (1993) Calcium plays a complex role in the regulation of melanogenesis in murine B16 melanoma cells. Pigment cell Res 6: 385–393

Cai C, Tu J, Najarro J, Zhang R, Fan H, Zhang FQ, Li J, Xie Z, Su R, Dong L, et al (2024) NRAS mutant dictates AHCYL1-governed ER calcium homeostasis for melanoma tumor growth. Mol Cancer Res 22: 386–401

Cárdenas C, Miller RA, Smith I, Bui T, Molgó J, Müller M, Vais H, Cheung KH, Yang J, Parker I, et al (2010) Essential Regulation of Cell Bioenergetics by Constitutive InsP3 Receptor Ca2+ Transfer to Mitochondria. Cell 142: 270–283

Cieri D, Vicario M, Giacomello M, Vallese F, Filadi R, Wagner T, Pozzan T, Pizzo P, Scorrano L, Brini M, et al (2017) SPLICS: a split green fluorescent protein-based contact site sensor for narrow and wide heterotypic organelle juxtaposition. Cell Death Differ 2018 *256* 25: 1131–1145

Daniele T, Hurbain I, Vago R, Casari G, Raposo G, Tacchetti C & Schiaffino MV (2014) Mitochondria and melanosomes establish physical contacts modulated by Mfn2 and involved in organelle biogenesis. 24: 393–403

Davis LC, Morgan AJ & Galione A (2020) NAADP -regulated two-pore channels drive phagocytosis through endo-lysosomal Ca 2+ nanodomains, calcineurin and dynamin. EMBO J 39

Fink BD, Bai F, Yu L & Sivitz WI (2017) Regulation of ATP production: Dependence on calcium concentration and respiratory state. Am J Physiol - Cell Physiol 313: C146–C153

Garrity AG, Wang W, Collier CMD, Levey SA, Gao Q & Xu H (2016) The endoplasmic reticulum, not the pH gradient, drives calcium refilling of lysosomes. Elife 5: 1–34

Grimm C, Jörs S, Guo Z, Obukhov AG & Heller S (2012) Constitutive Activity of TRPML2 and TRPML3 Channels versus Activation by Low Extracellular Sodium and Small Molecules. J Biol Chem 287: 22701

Hida T, Kamiya T, Kawakami A, Ogino J, Sohma H, Uhara H & Jimbow K (2020) Elucidation of Melanogenesis Cascade for Identifying Pathophysiology and Therapeutic Approach of Pigmentary Disorders and Melanoma. Int J Mol Sci 2020, Vol 21*, Page* 6129 21: 6129

Ho H & Ganesan AK (2011) The pleiotropic roles of autophagy regulators in melanogenesis. Pigment Cell Melanoma Res 24: 595–604

Hu Q-M, Yi W-J, Meng |, Su Y, Jiang S & Xu S-Z (2017) Induction of retinal-dependent calcium influx in human melanocytes by UVA or UVB radiation contributes to the stimulation of melanosome transfer. Cell Prolif 50: 50

Huang P, Zou Y, Zhong XZ, Cao Q, Zhao K, Zhu MX, Murrell-Lagnado R & Dong XP (2014) P2X4 Forms Functional ATP-activated Cation Channels on Lysosomal Membranes Regulated by Luminal pH. J Biol Chem 289: 17658–17667

Jia Q, Tian W, Li B, Chen W, Zhang W, Xie Y, Cheng N, Chen Q, Xiao J, Zhang Y, et al (2021) Transient Receptor Potential channels, TRPV1 and TRPA1 in melanocytes synergize UV-dependent and UV-independent melanogenesis. Br J Pharmacol 178: 4646–4662

Kaizuka T, Morishita H, Hama Y, Tsukamoto S, Matsui T, Toyota Y, Kodama A, Ishihara T, Mizushima T & Mizushima N (2016) An Autophagic Flux Probe that Releases an Internal Control. Mol Cell 64: 835–849

Katayama H, Kogure T, Mizushima N, Yoshimori T & Miyawaki A (2011) A Sensitive and Quantitative Technique for Detecting Autophagic Events Based on Lysosomal Delivery. Chem Biol 18: 1042–1052

Kim HRK, Lee GH, Bhattarai KR, Lee MS, Back SH, Kim HRK & Chae HJ (2021) TMBIM6 (transmembrane BAX inhibitor motif containing 6) enhances autophagy through regulation of lysosomal calcium. Autophagy 17: 761–778

Lee KW, Ryu KJ, Kim M, Lim S, Kim J, Kim JY, Hwangbo C, Yoo J, Cho YY & Kim KD (2024) RCHY1 and OPTN are required for melanophagy, selective autophagy of melanosomes. Proc Natl Acad Sci U S A 121: e2318039121

Li C, Wang X, Li X, Qiu K, Jiao F, Liu YY, Kong Q, Liu YY & Wu Y (2019) Proteasome Inhibition Activates Autophagy-Lysosome Pathway Associated With TFEB Dephosphorylation and Nuclear Translocation. Front Cell Dev Biol 7: 472905

Li M, Zhang WK, Benvin NM, Zhou X, Su D, Li H, Wang S, Michailidis IE, Tong L, Li X, et al (2017) Structural basis of dual Ca 2+ /pH regulation of the endolysosomal TRPML1 channel. Nat Struct Mol Biol 24: 205–213

Liu K, Zhang Z, Xu Y, Wu Y, Lian P, Ma Z, Tang Z, Zhang X, Yang X, Zhai H, et al (2024) AMPK-mediated autophagy pathway activation promotes ΔFosB degradation to improve levodopa-induced dyskinesia. Cell Signal 118: 111125

Lloyd-Evans E, Waller-Evans H, Peterneva K & Platt FM (2010) Endolysosomal calcium regulation and disease. Biochem Soc Trans 38: 1458–1464

Di Malta C, Cinque L & Settembre C (2019) Transcriptional regulation of autophagy: Mechanisms and diseases. Front Cell Dev Biol 7: 1–10

Mammucari C, Raffaello A, Vecellio Reane D, Gherardi G, De Mario A & Rizzuto R (2018) Mitochondrial calcium uptake in organ physiology: from molecular mechanism to animal models. Pflugers Arch 470: 1165–1179

Medina DL, Di Paola S, Peluso I, Armani A, De Stefani D, Venditti R, Montefusco S, Scotto-Rosato A, Prezioso C, Forrester A, et al (2015) Lysosomal calcium signalling regulates autophagy through calcineurin and TFEB. Nat Cell Biol 17: 288–299

Miao Y, Li G, Zhang X, Xu H & Abraham SN (2015) A TRP channel senses lysosome neutralization by pathogens to trigger their expulsion. Cell 161: 1306–1319

Motiani RK, Tanwar J, Raja DA, Vashisht A, Khanna S, Sharma S, Srivastava S, Sivasubbu S, Natarajan VT & Gokhale RS (2018) STIM1 activation of adenylyl cyclase 6 connects Ca 2+ and cAMP signaling during melanogenesis. EMBO J 37: e97597

Natarajan VT, Ganju P, Ramkumar A, Grover R & Gokhale RS (2014a) Multifaceted pathways protect human skin from UV radiation. Nat Chem Biol 10: 542–551

Natarajan VT, Ganju P, Singh A, Vijayan V, Kirty K, Yadav S, Puntambekar S, Bajaj S, Dani PP, Kar HK, et al (2014b) IFN-γ signaling maintains skin pigmentation homeostasis through regulation of melanosome maturation. Proc Natl Acad Sci U S A 111: 2301–2306

Palmieri M, Pal R, Nelvagal HR, Lotfi P, Stinnett GR, Seymour ML, Chaudhury A, Bajaj L, Bondar V V., Bremner L, et al (2017) MTORC1-independent TFEB activation via Akt inhibition promotes cellular clearance in neurodegenerative storage diseases. Nat Commun 8

Pan B, Li J, Parajuli N, Tian Z, Wu P, Lewno MT, Zou J, Wang W, Bedford L, Mayer RJ, et al (2020) The Calcineurin-TFEB-p62 Pathway Mediates the Activation of Cardiac Macroautophagy by Proteasomal Malfunction. Circ Res 127: 502–518

Park HJ, Jo DS, Choi H, Bae JE, Park NY, Kim JB, Choi JY, Kim YH, Oh GS, Chang JH, et al (2020) Melasolv induces melanosome autophagy to inhibit pigmentation in B16F1 cells. PLoS One 15: 1–10

Park K, Lim H, Kim J, Hwang Y, Lee YS, Bae SH, Kim H, Kim H, Kang SW, Kim JY, et al (2022) Lysosomal Ca2+-mediated TFEB activation modulates mitophagy and functional adaptation of pancreatic β-cells to metabolic stress. Nat Commun 13 doi:10.1038/s41467-022-28874-9 [PREPRINT]

Pathak T & Trebak M (2018) Mitochondrial Ca2+ signaling. Pharmacol Ther 192: 112–123

Prajapat SK, Mishra L, Khera S, Owusu SD, Ahuja K, Sharma P, Choudhary E, Chhabra S, Kumar N, Singh R, et al (2024) Methotrimeprazine is a neuroprotective antiviral in JEV infection via adaptive ER stress and autophagy. EMBO Mol Med 16: 185–217

Pryor PR, Reimann F, Gribble FM & Luzio JP (2006) Mucolipin-1 is a lysosomal membrane protein required for intracellular lactosylceramide traffic. Traffic 7: 1388–1398

Qu J, Yan M, Fang Y, Zhao J, Xu T, Liu F, Zhang K, He L, Jin L & Sun D (2023) Zebrafish in dermatology: a comprehensive review of their role in investigating abnormal skin pigmentation mechanisms. Front Physiol 14: 1296046

Roczniak-Ferguson A, Petit CS, Froehlich F, Qian S, Ky J, Angarola B, Walther TC & Ferguson SM (2012) The transcription factor TFEB links mTORC1 signaling to transcriptional control of lysosome homeostasis. Sci Signal 5

Ryu HY, Kim LE, Jeong H, Yeo BK, Lee JW, Nam H, Ha S, An HK, Park H, Jung S, et al (2021) GSK3B induces autophagy by phosphorylating ULK1. Exp Mol Med 2021 *533* 53: 369–383

Sarkar S, Floto RA, Berger Z, Imarisio S, Cordenier A, Pasco M, Cook LJ & Rubinsztein DC (2005) Lithium induces autophagy by inhibiting inositol monophosphatase. J Cell Biol 170: 1101–1111

Schmiege P, Fine M, Blobel G & Li X (2017) Human TRPML1 channel structures in open and closed conformations. Nat 2017 *5507676* 550: 366–370

Scotto Rosato A, Montefusco S, Soldati C, Di Paola S, Capuozzo A, Monfregola J, Polishchuk E, Amabile A, Grimm C, Lombardo A, et al (2019) TRPML1 links lysosomal calcium to autophagosome biogenesis through the activation of the CaMKKβ/VPS34 pathway. 10: 1–16

Sharma N, Sharma A & Motiani RK (2023) A novel gain of function mutation in TPC2 reiterates pH-pigmentation interplay: Emerging role of ionic homeostasis as a master pigmentation regulator. Cell Calcium 111: 102705

Shen D, Wang X, Li X, Zhang X, Yao Z, Dibble S, Dong XP, Yu T, Lieberman AP, Showalter HD, et al (2012) Lipid storage disorders block lysosomal trafficking by inhibiting a TRP channel and lysosomal calcium release. Nat Commun 2012 *31* 3: 1–11

Singh SK, Baker R, Sikkink SK, Nizard C, Schnebert S, Kurfurst R & Tobin DJ (2017) E-cadherin mediates ultraviolet radiation- and calcium-induced melanin transfer in human skin cells. Exp Dermatol 26: 1125–1133

Su H & Wang X (2020) Proteasome malfunction activates the PPP3/calcineurin-TFEB-SQSTM1/p62 pathway to induce macroautophagy in the heart. Autophagy 16: 2114–2116

Tanwar J, Ahuja K, Sharma A, Sehgal P, Ranjan G, Sultan F, Agrawal A, D’Angelo D, Priya A, Yenamandra VK, et al (2024) Mitochondrial calcium uptake orchestrates vertebrate pigmentation via transcriptional regulation of keratin filaments. 22: e3002895

Tanwar J, Saurav S, Basu R, Singh JB, Priya A, Dutta M, Santhanam U, Joshi M, Madison S, Singh A, et al (2022a) Mitofusin-2 Negatively Regulates Melanogenesis by Modulating Mitochondrial ROS Generation. Cells 11: 701

Tanwar J, Sharma A, Saurav S, Shyamveer, Jatana N & Motiani RK (2022b) MITF is a novel transcriptional regulator of the calcium sensor STIM1: Significance in physiological melanogenesis. J Biol Chem 298

Tanwar J, Singh JB & Motiani RK (2020) Molecular machinery regulating mitochondrial calcium levels: The nuts and bolts of mitochondrial calcium dynamics. Mitochondrion 57: 9–22

Toyoda M, Luo Y, Makino T, Matsui C & Morohashi M (1999) Calcitonin gene-related peptide upregulates melanogenesis and enhances melanocyte dendricity via induction of keratinocyte-derived melanotrophic factors. J Investig Dermatology Symp Proc 4: 116–125

Viollet B (2017) The energy sensor AMPK: Adaptations to exercise, nutritional and hormonal signals. Res Perspect Endocr Interact: 13–24

Wang C, Wang H, Zhang D, Luo W, Liu R, Xu D, Diao L, Liao L & Liu Z (2018) Phosphorylation of ULK1 affects autophagosome fusion and links chaperone-mediated autophagy to macroautophagy. Nat Commun 2018 91 9: 1–15

Wang J, Gong J, Wang Q, Tang T & Li W (2022a) VDAC1 negatively regulates melanogenesis through the Ca2+-calcineurin-CRTC1-MITF pathway. Life Sci Alliance 5

Wang S, Li H, Yuan M, Fan H & Cai Z (2022b) Role of AMPK in autophagy. Front Physiol 13: 1015500

Williams JA, Hou Y, Ni HM & Ding WX (2013) Role of Intracellular Calcium in Proteasome Inhibitor-induced Endoplasmic Reticulum Stress, Autophagy and Cell Death. Pharm Res 30: 2279

Xiong J & Zhu MX (2016) Regulation of lysosomal ion homeostasis by channels and transporters. Sci China Life Sci 59: 777–791

Xiong Q, Feng R, Fischer S, Karow M, Stumpf M, Meßling S, Nitz L, Müller S, Clemen CS, Song N, et al (2023) Proteasomes of Autophagy-Deficient Cells Exhibit Alterations in Regulatory Proteins and a Marked Reduction in Activity. Cells 12

Xu H, Delling M, Li L, Dong X & Clapham DE (2007) Activating mutation in a mucolipin transient receptor potential channel leads to melanocyte loss in varitint-waddler mice. Proc Natl Acad Sci U S A 104: 18321–18326

Xu Z, Liu Q, Li J, Wang J, Yang Z, Wang J, Gao L, Cheng J, He J, Dong Y, et al (2024) AMPKα is active in autophagy of endothelial cells in arsenic-induced vascular endothelial dysfunction by regulating mTORC1/p70S6K/ULK1. Chem Biol Interact 388: 110832

Yoshii SR & Mizushima N (2017) Monitoring and measuring autophagy. Int J Mol Sci 18: 1–13

Zajac M, Mukherjee S, Anees P, Oettinger D, Henn K, Srikumar J, Zou J, Saminathan A & Krishnan Y (2024) A mechanism of lysosomal calcium entry. Sci Adv 10

Zeng J, Acin-Perez R, Assali EA, Martin A, Brownstein AJ, Petcherski A, Fernández-del-Rio L, Xiao R, Lo CH, Shum M, et al (2023) Restoration of lysosomal acidification rescues autophagy and metabolic dysfunction in non-alcoholic fatty liver disease. Nat Commun 2023 *141* 14: 1–17

Zheng Q, Chen Y, Chen D, Zhao H, Feng Y, Meng Q, Zhao Y & Zhang H (2022) Calcium transients on the ER surface trigger liquid-liquid phase separation of FIP200 to specify autophagosome initiation sites. Cell 185: 4082–4098.e22

Zheng Q, Zhang H, Zhao H, Chen Y, Yang H, Li T, Cai Q, Chen Y, Wang Y, Zhang M, et al (2025) Ca2+/calmodulin-dependent protein kinase II β decodes ER Ca2+ transients to trigger autophagosome formation. Mol Cell 85: 620–637.e6

Zhu K, Dunner K & McConkey DJ (2010) Proteasome inhibitors activate autophagy as a cytoprotective response in human prostate cancer cells. Oncogene 29: 451–462

Zou J, Hu B, Arpag S, Yan Q, Hamilton A, Zeng YS, Vanoye CG & Li J (2015) Reactivation of Lysosomal Ca2+ Efflux Rescues Abnormal Lysosomal Storage in FIG4-Deficient Cells. J Neurosci 35: 6801–6812.

